# Nrf2 promotes thyroid development and hormone synthesis

**DOI:** 10.1101/2022.02.27.482168

**Authors:** Gillotay Pierre, Romitti Mirian, Dassy Benjamin, Haerlingen Benoit, Parakkal Shankar Meghna, Faria Fonseca Barbara, Ziros G. Panos, Pal Singh Sumeet, Sykiotis P. Gerasimos, Costagliola Sabine

## Abstract

In all vertebrates, the function of the thyroid gland is to capture iodide to synthesize hormones that act on almost all tissues and are essential for normal growth and metabolism. Thyroid hormone production is a multistep process that ultimately leads to the coupling of iodine to thyroglobulin, the matrix glycoprotein of hormone synthesis. This coupling is dependent on H2O2, the accumulation of which in thyroid tissue, a true iodine trap, causes a basal level of oxidative stress higher than in other tissues, which, if left unchecked, leads to cellular damage. The thyroid has efficient antioxidant and detoxifying enzymes that help it resist H202-induced oxidative stress and maintain the homeostasis necessary for hormone synthesis. By regulating the expression of genes involved in cellular detoxification processes, the transcription factor NRF2 acts as a master regulator of the cellular defense system against oxidative stress. Using zebrafish embryos and mouse thyroid organoids, we show here that direct control of thyroglobulin expression by NRF2 is an evolutionarily conserved chore mechanism in vertebrates to complete the necessary molecular defense arsenal against oxidative stress in thyroid tissue.

## Introduction

Protection against oxidative-stress induced damage is a common feature of every living organisms (Recently reviewed by R Siauciunaite and colleagues in 2019(Siauciunaite *et al*., 2019)) and is mediated by a wide range of biological process including (but not limited to) glutathione-induced gene expression (Hayes and McLellan, 1999), regulation of NADPH levels via the pentose phosphate pathways (Ralser *et al*., 2007) and the superoxide dismutases (Muid, Karakaya and Koc, 2014) and catalases (Chelikani, Fita and Loewen, 2004) activity. In addition, in many species, the nuclear erythroid factor 2 like 2 transcription factor (NRF2, encoded by the NFE2L2 gene in Humans) acts as a master-regulator of the cells oxidative and metabolic stress defence system (Mukaigasa *et al*., 2012; Loboda *et al*., 2016; Thanas *et al*., 2020). In physiological conditions, Nrf2 is sequestrated in the cytoplasm by its repressor the Kelch-like ECH-associated protein 1 (Keap1). To efficiently repress Nrf2, Keap1 forms a complex with Cul3 to recruit an E3 ubiquitin ligase complex that will drive the polyubiquitination and subsequent proteasomal degradation of Nrf2. Under stress conditions, the conformation of the Keap1-Cul3 complex will change, preventing further ubiquitination and degradation of newly formed Nrf2 protein, therefore allowing its translocation to the nucleus to regulate detoxifying and antioxidant genes expression (Iso *et al*., 2016; Yamamoto, Kensler and Motohashi, 2018; Renaud *et al*., 2019). To regulate gene expression, Nrf2 protein will bind to antioxidant response element (ARE) located in the promoter or enhancer region of target gene (Rushmore, Morton and Pickett, 1991; Zhu *et al*., 2016).

Because of its functional need of H_2_O_2_, the regulation of O.S in the thyroid tissue has attracted growing attention over the last few years (Renaud *et al*., 2019; Thanas *et al*., 2020). In all vertebrates, the thyroid gland plays a primordial role during the development and acts as a central regulator of the physiology of any individual (Yen *et al*., 2006; Mullur, Liu and Brent, 2014; Ortiga-Carvalho *et al*., 2016). During embryonic development, the thyroid enables the production of growth hormones (Shields *et al*., 2011; Liu *et al*., 2015) and plays a critical role in brain organogenesis and maturation (Mohan *et al*., 2012; Moog *et al*., 2017). In adults, it controls the basal metabolism, the cardiac function and the body temperatures among other functions (Klein and Ojamaa, 2001; Kahaly and Dillmann, 2005; Kim, 2008; Mullur, Liu and Brent, 2014; Ortiga-Carvalho *et al*., 2016). Congenital hypothyroidism (C.H.) is the most common congenital endocrine disorder (affecting 1 on every 3500 birth) and is characterised by a lack of physiological thyroid function at birth. This pathology can be caused by a defect affecting either the thyroid organogenesis or the thyroid hormones synthesis *per se* (Wassner, 2018). If left untreated, C.H. will cause severe mental and growth retardation in patient among other physiological consequences (Persani *et al*., 2018). Thyroid functions are mediated through both triiodothyronine (T3 - biologically active but produced to a lesser extend) and thyroxine (T4 - biologically less active but produced in high quantity) that are the main hormones it produces (Carvalho and Dupuy, 2017). Thyroid hormones (T.H.) production is an evolutionary conserved multi-step process ultimately leading to the H_2_O_2_-dependent coupling of iodine and thyroglobulin thereby creating iodinated-thyroglobulin, the precursor of the thyroid hormones (Di Jeso and Arvan, 2016; Carvalho and Dupuy, 2017). Consequently, as soon as the T.H. production machinery is functional, the thyroid gland undergoes a higher basal level of H_2_O_2_-induced oxidative stress (O.S) compared to other tissues.

Although a minimal level of O.S is mandatory for thyroid cells growth, function and proliferation (Poncin, Colin and Gérard, 2009; Poncin *et al*., 2010), if left unchecked, higher level of H_2_O_2_ will break the balance between oxidant product and thyroid’s antioxidant defences, ultimately resulting in O.S-induced damage that can lead to genomic and epigenetic mutation (El Hassani *et al*., 2019; Renaud *et al*., 2019).

Recent studies performed on adult mice thyroid glands demonstrated the role of Nrf2 as a direct controller of the thyroglobulin (Tg) expression and as a central actor of the thyroid gland stress defense system (Renaud *et al*., 2019; Chartoumpekis *et al*., 2020) Despite these important studies related with oxidative stress in mature thyroid follicular cells, we lack information on the potential role of *Nrf2* during thyroid development. Here, using zebrafish embryos and mouse-embryonic stell cell (mESC)-derived thyroid follicles, we though to characterise the role of *Nrf2/nrf2a* during mammalian and non-mammalian thyroid development.

## Results

### Identification of *nrf2a* as an actor of zebrafish thyroid functional maturation

Due to whole genome duplication, zebrafish genome encodes for two orthologues of mammalian Nrf2: *nrf2a* (encoded by the *nfe2l2a* gene, ZDB-GENE-030723-2) and *nrf2b* (encoded by the *nfe2l2b* gene, ZDB-GENE-120320-3) both encoding for transcription factors (Hahn *et al*., 2011). To assess whether one of these two paralogues plays a role during thyroid development or thyroid functional maturation, we generated single guide-RNA (sgRNA) targeting the DNA binding domain of *nrf2a* and *nrf2b*, respectively located on their exon 5 and 4. To uncover potential thyroid developmental defects, we took advantage of our previously published F0 Crispr/Cas9-based screening approach (Trubiroha *et al*., 2018). Here, following one-cell stage injection in Tg(*tg:nlsEGFP*) embryos (Trubiroha *et al*., 2018), thyroid development was monitored *in vivo* at 55, 72 and 144 hours post fertilisation (hpf) for putative developmental defect affecting either early thyroid development, proliferation and migration or thyroid functional maturation respectively. In these experiments two control were used: 1) the injection of a sgRNA targeting *adamtsl2* as a control for false positive phenotypes (Trubiroha *et al*., 2018) and 2) analysis and raising of non-injected embryos as a fertility and development quality control. At 55 and 80hpf, the thyroid development of the nrf2a and nrf2b injected embryos were similar and no gross developmental defect were observed compared to their *adamtsl2* crispants and non-injected siblings (Sup Fig. 1 A-H). However, at 144hpf, thyroid tissue appeared enlarged in around 55% (170/312) of the *nrf2a* crispants compared to their *adamtsl2* crispants and non-injected siblings (Sup Fig. 1 I-L). Conversely, we could not uncover any gross or thyroid-related developmental defect in *nrf2b* F0 crispants compared to their *adamtsl2* crispants and non-injected siblings despite high mutagenic activity of the sgRNA used in these experiments (data not shown). Considering the reproducibility of the observed phenotype in F0, we decided to generate a stable nrf2a mutant line to further investigate the mechanism underlying the observed developmental defect of the thyroid gland.

### Generation of a stable *nrf2a* mutant line

To assess the transmission of the mutated alleles and associated thyroid phenotype, we raised F0 Tg(*tg:nlsEGFP*) *nrf2a* crispants to adulthood and outcrossed them with AB* wild-type (WT) animals in order to identify and select transmitted mutated alleles. F1 progeny was raised to adulthood and subsequently genotyped to characterise the transmitted mutated alleles using Sanger sequencing. We identified several mutated *nrf2a* alleles harbouring insertion, deletion or a mix of both and selected an allele presenting a deletion of 5 nucleotides between the nucleotides 1450 and 1454 of the reference coding sequence, this deletion is located in the exon 5 prior to the DNA binding domain of the Nrf2a protein (here after referenced as *nrf2aΔ5*) (Fig. 1A). The *nrf2aΔ5* allele is predicted to induce a frameshift leading to a truncated Nrf2a protein of 550 amino acids (aa), while the WT protein is 587 amino acids long, including 67 incorrect amino acids at the C-terminus. *In silico* analysis of the mutated protein encoded by the *nrf2aΔ5* allele revealed that, out of the 112 amino acids composing the WT protein coiled-coil domain in which the DNA binding region resides, only 8 amino acids were conserved in the mutated protein, suggesting that it is non-functional. To experimentally assess the functional consequences of our *nrf2aΔ5* mutated allele, we tested its ability to activate the transcription of an ARE-driven firefly luciferase reporter gene following a previously published protocol by Mukaigasa and colleagues in 2012 (Mukaigasa *et al*., 2012). In these experiments, comparison was made with wild-type Nrf2a protein and the previously published Nrf2a fh318 mutated protein known to be non-functional (Mukaigasa *et al*., 2012). After transient transfection in HEK cells, WT Nrf2a protein strongly activates the transcription of the firefly luciferase reporter gene whereas both the *nrf2aΔ5* and *nrf2a* fh318 mutants were able to induce only weak expression of the reporter gene, if at all. (Fig. 1B). This shows that our *nrf2aΔ5* allele, like the *nrf2a* fh318 mutant alleles, encodes a nonfunctional transcription factor.

**Figure 1.**
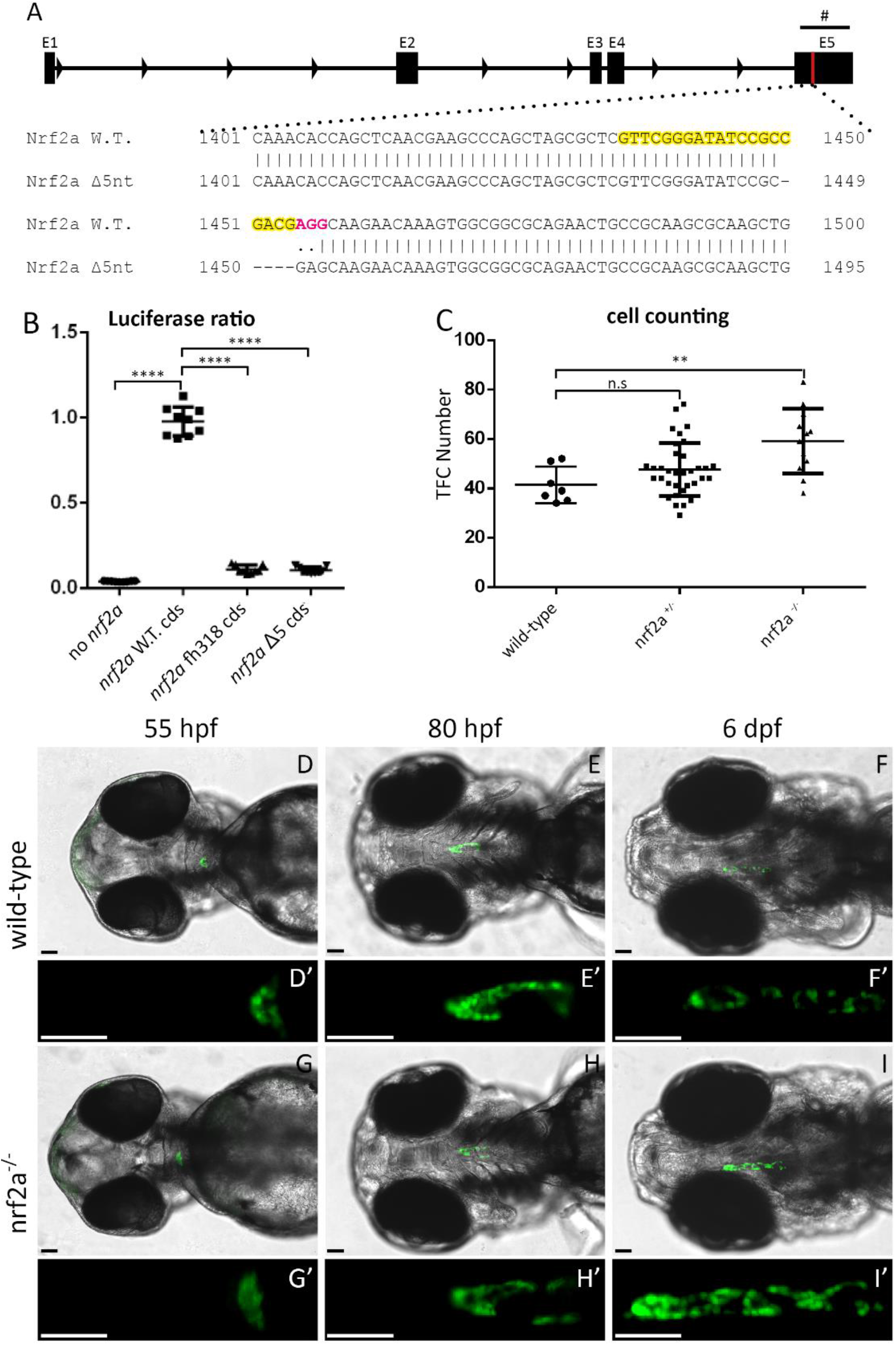
Live imaging of transgenic Tg(tg:nlsEGFP) zebrafish embryos carrying a 5nt deletion in *nrf2a* allows real-time in vivo analysis of thyroid development. (A) A 5nt deletion was generated in the exon 5 (E5) of the nrf2a gene resulting in a frame-shift inducing premature stop codon disturbing the DNA binding domain of the gene. Guide sequence highlighted in yellow with PAM sequence highlighted in magenta. (B) Activation of an ARE-driven luciferase reporter by WT and mutated nrf2a. HEK cells were transfected with plasmids encoding for the full-length coding sequences of a WT nrf2a, nrf2a fh318 or nrf2a D5 to evaluate their respective ability to drive luciferase expression. Results show that WT nrf2a (second from the left) is able to drive expression of luciferase while *nrf2a* fh318 and Δ5 are unable to drive luciferase expression (third and fourth from the left respectively). Results are shown as mean +/− SD. Asterisk denotes significant differences between positive control (WT nrf2a cds) and test conditions (**** p <0.0001, Dunnett’s multiple comparison test) (C) Quantification of thyroid follicular cell (TFC) number in individual wild-type (n=7), heterozygous (n=34) and homozygous mutant embryos (n=12) show significant difference between wild-type and homozygous mutant. Results are shown in mean +/− SD. Asterisk denotes significant differences between groups of embryos (** P < 0.01, Kruskal-Wallis multiple comparison test). (D-I) show pictures of the whole head of a representative embryo at specified developmental stage (ventral view, anterior is to the left). (D’-I’) show an enlarged picture of the thyroid region of the corresponding embryo. Epifluorescence microscopy of live Tg(tg:nlsEGFP) zebrafish reveals an increase in the thyroid size at 6 days post fertilisation (dpf) in the mutant embryos compared to their wild-type and heterozygous (data not shown) siblings. Live-imaging also reveals that prior to 4dpf thyroid development shows no apparent differences between mutant, heterozygous (data not shown) and wild-type embryos. Scale bars: 50μm (D-I’). “#” indicates de DNA binding domain of the *nrf2a* gene.

### *nrf2a* loss-of-function induces hypothyroidism in homozygous mutant zebrafish embryos

We then analysed, *in vivo*, the thyroid phenotype of Tg(*tg:EGFPnls*) *nrf2aΔ5* homozygous embryos to see if they exhibited thyroid defects similar to those observed in F0 crispants. Similar to what we observed previously in crispants, thyroid development at 55 (Fig. 1D and G) and 80 hpf (Fig. 1E and H) in *nrf2aΔ5* hemi- and homozygous embryos does not differ from their WT siblings. However, at 144 hpf, the thyroid gland of *nrf2aΔ5* homozygous embryos appears enlarged along the antero-posterior axis compared with the gland of their hemizygous mutant and WT siblings (Fig. 1F and I). To quantify this apparent thyroid enlargement, we used confocal live imaging on progeny derived from an incross of Tg(*tg:nlsEGFP*); *nrf2aΔ5* hemizygous adult animals to perform thyroid cell counting at 144 hpf (Fig. 1C). On average, thyroid glands of WT embryos were composed of 41.4 +/− 7.3 cells while thyroids of homozygous *nrf2aΔ5* embryos were composed of 59.2 +/− 13.1 cells, showing an enlargement of approximately 40% of their thyroid gland. Excessive proliferation of the thyroid follicular cells (TFC) is often associated with continuous stimulation of the thyroid gland by the TSH signalling pathway. Such continuous stimulation may result from a defect in thyroid hormones (T.H) production, thus triggering the feedback loop controlling the hypothalamo-pituitary-thyroid axis (Dumont, Maenhaut and Lamy, 1992; Ortiga-Carvalho *et al*., 2016). To test this hypothesis, we performed whole mount immunofluorescence (WIF) for thyroxine (T4) and its precursor the iodinate-thyroglobulin (Tg-I) on 6 days post-fertilisation (dpf) progeny from *nrf2aΔ5* heterozygous carrier (Fig 2A-O). WIF staining for T4 and Tg-I confirmed that the thyroid enlargement observed in *nrf2aΔ5* homozygous coincides with dyshormonogenesis as shown by the lack of or barely detectable T4 and Tg-I staining (6.75*10^5^ +/− 2.68*10^5^ A.U for T4 and 1.36*10^6^ +/+ 6.61*10^5^ A.U for TG-I) compared with their WT siblings (2.10*10^6^ +/− 6.13*10^5^ A.U for T4 and 3.37*10^6^ +/−1.50*10^6^ A.U for TG-I) (Fig. 2Q and R). To ensure that lack of detectable colloidal T4 was not caused by a defect in thyroid follicle cell polarisation, we analysed the thyroid follicles by confocal imaging after immunostaining for ZO-1 protein which labels the apical junctions between the TFC. Imaging of WT and homozygous *nrf2aΔ5* embryos revealed no discernible differences in follicular polarisation between embryos (Supp Fig 2), suggesting that the observed deficiency of T4 production is not caused by a defect in thyroid follicle morphogenesis *per se*.

**Figure 2:**
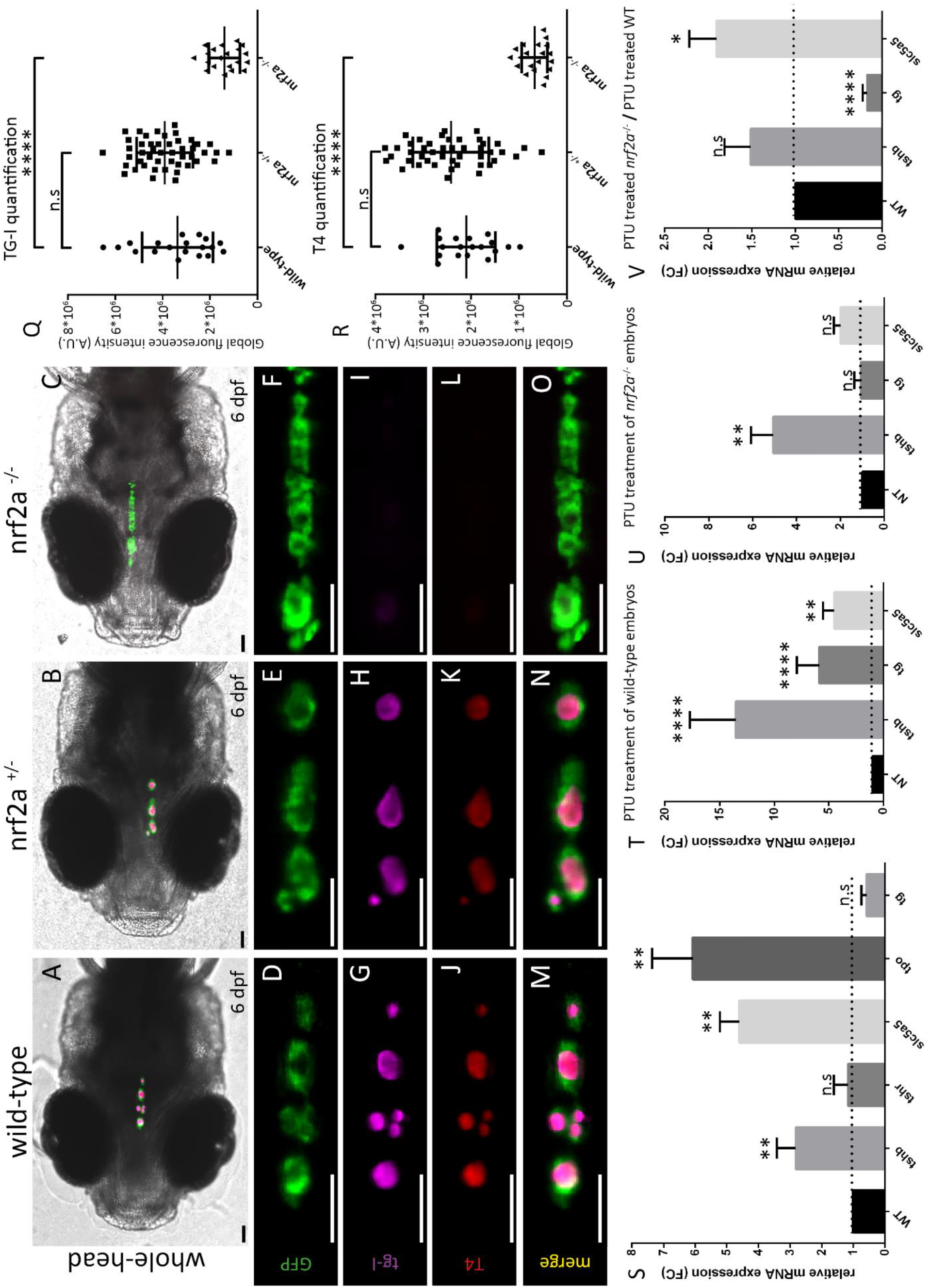
Loss-of-function of *nrf2a* induces thyroid dyshormonogenesis. (A-O) immunofluorescence on 6dpf Tg(tg:nlsEGFP) zebrafish embryos targeting GFP, thyroxine (T4 - Cy3 Red) and iodinated-thyroglobulin (TG-I - Cy5 magenta) reveals a defect in thyroid hormones production in *nrf2a* homozygous mutant compared to their wild-type and heterozygous mutant siblings. (A-C) show whole head pictures of a representative embryos of each genotype (ventral view anterior to the left). (D-O) enlarged views of the thyroid of the corresponding embryos demonstrated thyroid follicular organization in all embryos (D-F), however absence of TG-I and T4 was observed only in *nrf2a*^−/−^ embryos (F,I,L,O). Quantification of iodinated-thyroglobulin (Q) and colloidal thyroxine (R) immunofluorescence signal in individual wild-type (n=19), heterozygous (n=45) and homozygous mutant embryos (n=20) reveals significant reduction in *nrf2a*^−/−^ group. Results are shown in mean +/− SD. Asterisk denotes significant differences between groups of embryos (**** P < 0.0001, one way Anova multiple comparison test). Scale bars (A-O) 50 μm. (S) Quantitative analysis of thyroid markers expression levels by qPCR showed no differential expression of *tshr* (p value = 0.7706) and indicates a slight reduction, although not significant, of *tg* expression (p value = 0.9125) between wild-type and homozygous mutant embryos at 6dpf. However, mRNA levels show significant differences for *tsh*-β (p value < 0.01), *slc5a5* (p value < 0.01) and *tpo* (p value < 0.01). “n.s” denotes no significant difference between groups of embryos. (T-U) Quantitative analysis of thyroid markers expression levels of 6dpf wild-type and nrf2a-/- embryos following PTU treatment shows strong increase of *tshϐ* (p value < 0.0001), *tg* (p value < 0.0001) and *slc5a5* (p value < 0.01) in wild-type embryos and also shows an increase of *tshϐ* expression (p value < 0.01) in *nrf2a*^−/−^ embryos. However, analysis of the expression of *tg* (p value > 0.9999) and *slc5a5* (p value = 0.2499) in *nrf2a*^−/−^ showed no differences between PTU-treated and non-treated embryos. (V) Comparative analysis of the effect of PTU treatment between *nrf2a*^−/−^ embryos and their wild-type siblings. Quantitative analysis showed that the expression of *tsh*b does not significantly change between *nrf2a*^−/−^ and their wild-type siblings (p value > 0.9999). However, analysis revealed that nrf2a-/- embryos are not able to over-express *tg* upon PTU treatment (p value < 0.0001) while the increase of *slc5a5* expression is higher in *nrf2a*^−/−^ treated embryos (p value = 0.0183) compared to their treated wild-type siblings. All qPCR data are represented as mean +/− SD. Asterisk denotes significant difference between groups of embryos (* p < 0.05, ** p < 0.01 and **** p < 0.0001, Kruskal-Wallis multiple comparison test). “n.s” denotes no significant difference between groups of embryos. N varies between 5 and 6 individual pool of 10 embryos for each gene.

### Loss *of nrf2a* function induces a down regulation of *tg* expression

The phenotype of thyroid dyshormonogenesis observed in homozygous nrf2aΔ5 mutants may be due to several causes. Indeed, T.H. synthesis requires a complex machinery allowing uptake and transport of iodide from the bloodstream to the colloidal space, coupling of iodine to the thyroglobulin and eventually proteolysis of iodinated-thyroglobulin and release of T.H. in the bloodstream (Carvalho and Dupuy, 2017). As immunostaining for T4 and Tg-I reveals an absence of both signal in *nrf2aΔ5* homozygous mutants, we decided to analyse the expression of thyroid functional markers and stimulating factors by qPCR on 6dpf *nrf2aΔ5* zebrafish embryos (Fig. 2S). We first looked at the thyroid stimulating signalling by analysing the expression of *tsh-ϐ* and *tshr*. In *nrf2aΔ5* homozygous mutants, expression of *tsh-ϐ* was greatly increased compared with their WT siblings (Fig. 2S, 2^nd^ from the left), consistent with the known effects of a lack of T.H production (Shupnik *et al*., 1985; Chiamolera and Wondisford, 2009). On the other hand, qPCR experiments did not show any difference in the expression of *tshr* in *nrf2aΔ5* homozygous mutant embryos compared to their WT siblings (Fig. 2S, 3^rd^ from the left). As the absence of Tg-I staining in *nrf2aΔ5* homozygous mutant embryos seems to indicate a defect in the very first step of T.H. synthesis, we decided to analyse the expression levels of *slc5a5, tpo* and *tg* as these genes are critical component to the formation of iodinated-thyroglobulin (Carvalho and Dupuy, 2017). Consistent with the increase of *tshϐ* expression (Opitz *et al*., 2011), the expression levels of *slc5a5* and *tpo* were greatly increase in *nrf2aΔ5* homozygous mutants compared to their WT siblings (Fig. 2S, 4^th^ and 5^th^ from the left). However, the expression of *tg*, although not statistically significant, appeared reduced in *nrf2aΔ5* homozygous mutants compared with their wild-type siblings despite the increase of their TFC number (Fig. 2S, 6^th^ from the left). To confirm that the lack of *tg* upregulation was caused by defective *nrf2a* and not by an insufficient Tsh stimulation, we treated *nrf2aΔ5* homozygous mutant and wild-type embryos with 1-phenyl 2-thiourea (PTU), a chemical commonly used to inhibit pigmentation but shown to inhibit T.H production (Elsalini and Rohr, 2003; Opitz *et al*., 2011). We treated wild-type and *nrf2aΔ5* homozygous mutant embryos with PTU at a concentration of 30μg/mL from 24 to 144 hpf and performed qPCRs analysis for *tsh-ϐ, slc5a5* and *tg* at 144hpf. Upon PTU treatment, *tsh-ϐ* and *slc5a5a* expression were increased in both WT and *nrf2aΔ5* homozygous mutants compared to their nontreated siblings (Fig. 2T and U, 2^nd^ and 4^th^ from the left), confirming the efficiency of our treatment to inhibit T.H production and to trigger the Tsh-mediated feedback loop. Surprisingly, while PTU treatment induced an increase of *tg* expression in WT embryos (Fig. 2T, 3^rd^ from left), *tg* expression remained unaffected in *nrf2aΔ5* homozygous mutant embryos compared to their non-treated siblings (Fig. 2U, 3^rd^ from the left). Interestingly, when we compared the modulation of *tg* expression in both PTU-treated WT and *nrf2aΔ5* homozygous mutant embryos, we saw a marked reduction in *tg* expression in *nrf2aΔ5* homozygous mutant embryos (Fig. 2V, 3^rd^ from left). These results suggest that the lack of functional *nrf2a* affects the modulation of *tg* expression in zebrafish thyroid.

### The thyroid dyshormonogenesis observed in *nrf2a*Δ5 homozygous mutant embryos is a cell autonomous phenotype

Although the initial characterisation of the *nrf2aΔ5* mutant phenotype seems to indicate a role for *nrf2a* during thyroid functional maturation, we could not rule out the possibility that this phenotype is secondary to another defect affecting the whole zebrafish embryos. To exclude non-cell-autonomous effects, we sought to rescue the thyroid function by generating a new transgenic line Tg(*tg:nrf2a^T2A_mKO2nls^*) in which the thyroglobulin promoter drives the expression of a functional *nrf2a* coding sequence specifically in the thyroid follicular cells. Further, for visualization, the nrf2a coding sequence is linked, via a T2A viral peptide, to the nuclear localised monomeric Kusabira Orange 2 fluorescent protein (here after referred as the *nrf2a_mKO2* line or “rescue” allele). Although fluorescent reporter allows easy selection of transgenic embryos, we wanted to confirm whether the over-expression of *nrf2a* mRNA could be detected in the thyroid of transgenic embryos by WISH. Indeed, due to its short half-life (McMahon *et al*., 2004), detection of *nrf2a* mRNA is some tissue falls below the sensitivity of WISH staining. Following WISH staining, *nrf2a* can be easily detected in tissue such as the developing gills and the liver but remains undetectable in the developing thyroid in 72hpf WT embryos. By contrast, in addition to the other tissue, *nrf2a* can be detected in the thyroid of 72hpf *nrf2a_mKO2* embryos confirming that our transgenic line displays a thyroidal over-expression of *nrf2a* (Supp Fig 3).

**Figure 3:**
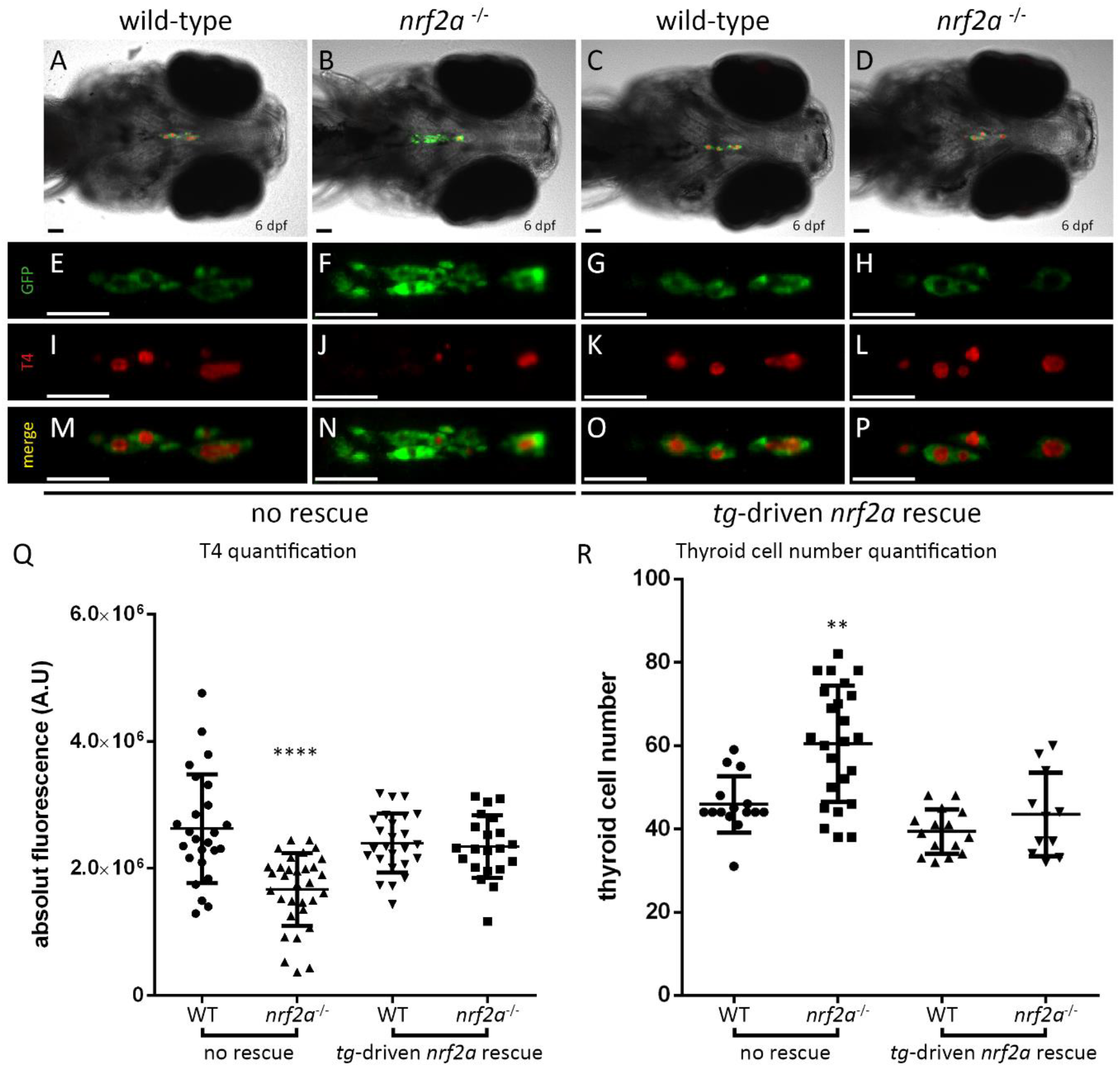
Thyroid-specific rescue of nrf2a function is sufficient to recover physiological T4 production. Immunofluorescence on 6dpf Tg(*tg:nlsEGFP; tg:nrf2a^T2A_mKO2^*) zebrafish embryos targeting GFP, and thyroxine (T4 - Cy3 Red) reveals a than the presence of the *tg:nrf2a*^T2A_mKO2^ construct is sufficient to rescue the thyroid hormone production defect in *nrf2a* homozygous mutants compared to their wildtype and heterozygous mutant siblings (A-P). (A-D) show whole head pictures of a representative embryos of each genotype (ventral view anterior to the left). (E-P) enlarged views of the thyroid of the corresponding embryos demonstrates thyroid follicular organization in all embryos (E-H) and reveals that the absence of T4 staining observed in nrf2a homozygous mutant embryos (second column to the left) is rescued in nrf2a homozygous mutant embryos expressing de *tg:nrf2a*^T2A_mKO2^ transgene (last column to the left). Scale bars: 50 μm. (Q) Quantification of colloidal thyroxine immunofluorescence signal in individual wild-type (n=25), *nrf2a* homozygous mutant embryos (n=32), WT *nrf2a^T2A_mKO2^* (n=25) and *nrf2a* homozygous mutant *nrf2a^T2A_mKO^2* embryos (n=21) confirmed the rescue of the T4 production in *nrf2a* homozygous mutant *nrf2a^T2A_mKO2^* embryos group compares to their siblings from other genotypes. Results are shown in mean +/− SD. Asterisk denotes significant differences between groups of embryos (**** p value < 0.0001, one way Anova multiple comparison test). (R) Quantification of thyroid follicular cell (TFC) number in individual wild-type (n=15) and homozygous *nrf2a* mutant embryos (n=24) and WT (n=15) and homozygous *nrf2a* mutant (n=11) *nrf2a*^T2A_mKO2^ embryos show that the *nrf2a*^T2A_mKO2^ transgene rescued the thyroid cell number in homozygous *nrf2a* mutant embryos. Results are shown in mean +/− SD. Asterisk denotes significant differences between groups of embryos (** p value < 0.01, Kruskal-Wallis multiple comparison test).

Using this transgenic line we generated animals carrying the rescue allele and the *nrf2aΔ5* mutation. We then assessed the thyroid function of the rescued embryos at 6dpf by T4 immunostaining as we had done previously (Fig. 3A-P). In wild-type animals, quantification of the T4 fluorescence signal showed that the presence of the rescue transgene within the thyroid cells did not significantly affect the ability of the thyroid to produce T4 (2,62*10^6^ +/−8.55*10^5^ A.U. in WT embryos vs 2.40*10^6^ +/− 4.64*10^5^ A.U. in WT *nrf2a_mKO2* embryos) (Fig. 3Q, 1^st^ and 2^nd^ from the left). However, in 6dpf *nrf2aΔ5* homozygous embryos, we could observe that T4 production was restored to physiological levels upon expression of the rescue allele (2.34*10^6^ +/− 4.93*10^5^ A.U. in rescued *nrf2aΔ5* homozygous embryos; 2,62*10^6^ +/− 8.55*10^5^ A.U. in WT embryos and 1.67*10^6^ +/− 5.72*10^5^ A.U. in non-rescued *nrf2aΔ5* homozygous embryos) (Fig 3Q, 3^rd^ and 4^th^ from left). In addition, in the presence of the rescue allele, we observed that the thyroid of *nrf2aΔ5* homozygous mutant embryos appeared to be of comparable size to that of WT *nrf2a_mKO2* embryos. To confirm this, we performed cell counting following confocal live imaging on 6dpf *nrf2aΔ5* and WT embryos carrying or not the rescue allele. The quantification of thyroid cell number revealed that the presence of the rescue allele in WT embryos does not significatively affect the thyroid cell number (Fig 3R, 1^st^ and 3^rd^ from the left). However, while the thyroid glands of *nrf2aΔ5* homozygous embryos were, on average, composed of 60.4 +/− 13.9 cells, they are composed of 43.4 +/− 10 cells when homozygous mutant embryos carry the rescue allele (Fig 3R, 2^nd^ and 4^th^ from left). Altogether these data support the cell-autonomous nature of thyroid dyshormonogenesis phenotype observed in *nrf2aΔ5* homozygous mutant zebrafish embryos.

Using zebrafish embryos, we showed that impairing the function of *nrf2a* strongly affects the thyroid functional maturation, leading to a lack of T.H production. We also proved that such defect does not results from a disorganisation of the thyroid follicular structure or from a lack of expression of genes involved in the T.H production machinery such as *slc5a5* and *tpo*, but rather from a reduced ability of the *nrf2aΔ5* homozygous mutant embryos to produce the thyroglobulin.

### Loss of *Nrf2* impairs differentiation of mESC-derived thyroid follicles

Recently, the role of *Nrf2* in thyroid gland physiology has been investigated by P. Ziros and colleagues using ubiquitous and thyroid-specific *Nrf2* Knock-out mouse model (Ziros *et al*., 2018). In this study, the authors described how loss of *Nrf2* function causes a thyroid phenotype only under stress conditions such as iodine overload. Thereby demonstrating that *Nrf2* plays a critical role in the regulation of oxidative and metabolic stress in adult thyroid gland. However, in the same study, the authors described that the loss of *Nrf2* function causes a reduction of the thyroglobulin expression in PCCL3 rat thyroid follicular cell culture. Importantly, this effect was observed without the need to induce stress condition in *Nrf2* KO PCCL3 cells (Ziros *et al*., 2018). Based on these results, we hypothesized that adult *Nrf2* KO mice might develop body-wide resistance to the effects of *Nrf2* defficiency which in turn, might reduce the visible effects on thyroid development and physiology. Therefore, to explore this aspect, we decided to use the *in vitro* model of mESC-derived thyroid follicles that we previously developed (Antonica *et al*., 2012; Romitti *et al*., 2021). mESC-derived thyroid follicles offer the unique advantage of recapitulating thyroid development in an ex *vivo* controlled medium which will help us characterise the effects of loss of *Nrf2* function on thyroid development and functional maturation.

The mouse orthologue of the zebrafish *nrf2a* is encoded by the *Nfe2l2* gene (also known as *Nrf2*) and is composed of five exons. Similarly to the zebrafish *nrf2a*, the DNA binding domain of the Nrf2 protein is located within the fifth exon. Thus, to generate a loss of function *Nrf2* similar to the zebrafish mutant line we generated, we targeted a region immediately upstream of the DNA binding domain using Crispr/Cas9 technology.

Following targeted mutagenesis, we identified two clones carrying mutations that lead to disruption of the DNA binding domain and to appearance of a premature STOP codon. The first clone (here after referred as “clone 16” or *Nrf2KO*, see Fig. 4A) harbours a deletion of 19 nucleotides and the second clone (here after referred as “clone 19”) harbours a deletion of one nucleotide.

**Figure 4:**
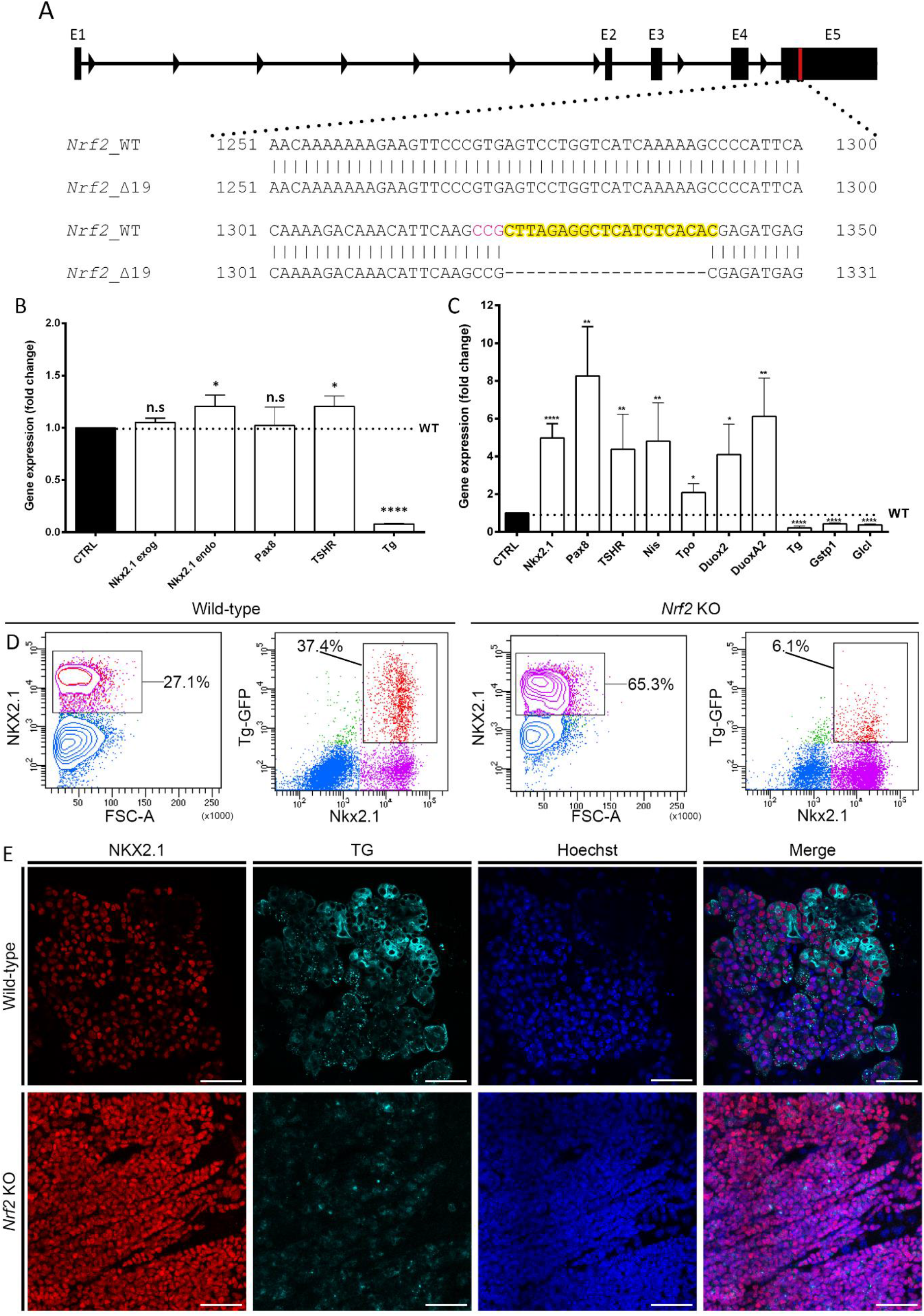
Analysis of Nrf2 KO mESC-derived thyroid organoids highlight the importance of Nrf2 during mammalian thyroid development. (A) A 19nt deletion was generated in the exon 5 (E5) of the Nrf2 gene resulting in a frame-shift inducing premature stop codon disturbing the DNA binding domain of the gene. Guide sequence highlighted in yellow with PAM sequence highlighted in magenta. (B) qPCR analysis performed at day 7 of the differentiation protocol demonstrates significant downregulation in Tg gene expression and slight upregulation in Nkx2.1 and Tshr gene expression. (C) qPCR analysis performed at the end of the differentiation protocol (at day 22) demonstrates significant up regulation in Nkx2.1, Pax8, Tshr, Nis, Tpo, Duox2 and Duoxa2 gene expression and significant downregulation in Tg, Gstp1 and Gclc gene expression. All statistical analysis performed using Mann-Whitney test, asterisks denote a significant difference compared to the wild-type cell line: *p<0.05, **p<0.01 and ****p<0.001. (D) Flow cytometry analysis of Wild-type (1st and 2nd panels from the left) and Nrf2 KO (3rd and 4th panels from the left) thyroid organoids after 22 days of differentiation. 1st and 3rd panel show the proportion of NKX2.1+ cells in the whole sample, percentage of NKX2.1+ cells are written. 2nd and 4th panel show the proportion of bTG+ cells within the Nkx2.1+ cell population, percentage of bTG+, NKX2.1+ cells are written. (E) Confocal images obtained following immunofluorescence experiments on fixed sample after 22 days of differentiation. Samples were marked for NKX2.1 (Red) and TG (Cyan). Nuclei were labelled using Hoechst (Blue).

We firstly validated the ability of both clones to overexpress Nkx2.1 and Pax8 following dox treatment at the 7^th^ day of the differentiation protocol using immuno-staining experiments (data not shown), suggesting that the loss of function of *Nrf2* does not affect the tetracyclin-inducible over-expression of *Nkx2.1* and *Pax8*.

We then differentiated both clones into thyroid organoids following our previously published protocol (Antonica *et al*., 2012; Romitti *et al*., 2021) to assess their ability to form thyroid follicles at the end of the protocol (day 22). Immunofluorescence analysis of fixed samples at day 22 were performed to analyse expression of thyroid markers (Nkx2.1 and Tg). Although both *Nrf2* KO clones can express Nkx2.1, they were unable to express Tg (Supp Fig. 4) and to organise into follicles. Considering that both clones presented similar phenotype we decided to focus our investigations using clone 16, therefore all following data and mention of the *Nrf2* KO refer to clone 16 cell line. Analysis of this mutation revealed that it leads to the apprearance of a premature stop codon resulting in a truncated protein of 469aa length, of which only the first 441aa are conserved with the 597aa long WT protein.

To better characterise the phenotype observed upon loss of *Nrf2* function, we analysed the expression of several thyroid markers at day 7 of our differentiation protocol. While *Nrf2* KO cells expressed similar levels of *Pax8* and slightly increased levels of *Tshr* and exogenous and endogenous *Nkx2.1* (Fig. 4B) compared with the control cell-line,, they showed significant downregulation of *Tg* expression (Fig 4B). On day 22 of the differentiation protocol, qPCR analysis revealed that Nrf2 KO cells expressed more *Nkx2.1, Pax8, Tshr, Nis* and *Tpo* and less *Tg*, similar to zebrafish embryos. In addition, we observed higher expression of *Duox2* and *Duoxa2* (Fig. 4C). Interestingly, we also observed a decrease in the expression of *Gstp1* and *Gclc* (Fig. 4C), two classical targets of Nrf2 (Mukaigasa *et al*., 2012; Renaud *et al*., 2019) confirming that our generated deletion of 19 nucleotides abrogates Nrf2 function. Analysis of the initial Nkx2.1 immunofluorescence experiments suggested that a higher proportion of cells in the Nrf2 KO cell culture were Nkx2.1+. To confirm this observation, we decided to perform flow cytometry analysis after Nkx2.1 immuno-staining and taking advantage of our Tg reporter (bTg-eGFP+). Flow cytometry quantification demonstrated higher percentage of Nkx2-1+ cells from the Nrf2 KO line (65.3%) compared with the control cell line (27.1%). Moreover, we observed that the percentage bTg-eGFP+ cells among Nrf2 KO-NKX2-1+ cells was significantly lower with only 6.1% of cells expressing eGFP while 37.4% expressed eGFP in the control cell line (Fig. 4D).

Immunofluorescence analysis of the ability of our Nrf2 KO cell line to form thyroid follicles revealed a lower number of follicular organised structures, defined by the presence of a lumen, than in the control cell line. Moreover, these experiments confirmed the decreased Tg levels in Nrf2 KO-differentiated cells, while the percentage of Nkx2-1 cells was higher compared with the control cell line. (Fig. 4D 1st and 3rd from the left).

### Functional Nrf2 is essential for thyroid function in mESC-derived follicles

One of the most important aspects of the *nrf2a* loss of function phenotype in zebrafish is the absence of thyroid hormones and iodinated-thyroglobulin production. To assess the conservation of this phenotype in our organoids, we analysed the presence of Nis at the basal membrane of follicles and the ability of Nrf2 KO cells to produce iodinated thyroglobulin. After 22 days of differentiation, we observed that Nis protein is well expressed at the membrane of both Nrf2 KO and control cell lines. However, the presence of Tg-I is almost exclusively detected in the lumen of *Nrf2* WT-derived follicles (Fig. 5A). Quantification analysis using iodine organification assays (Antonica *et al*., 2017) confirmed such observations. Indeed, while both cell lines are able to uptake iodine with the same efficiency (Fig 5B), the levels of protein-bound ^125^I were significantly lower in *Nrf2* KO thyroid cells (Fig. 5C). This results in a lower proportion of cells able to promote iodide organification in *Nrf2* KO-derived thyrocytes (3.43%) compared to control condition (22.45%) (Fig. 5D). Interestingly, as in the *nrf2aΔ5* zebrafish mutants, lack of Tg appears to be the primary cause of this defect in thyroid hormone production.

**Figure 5:**
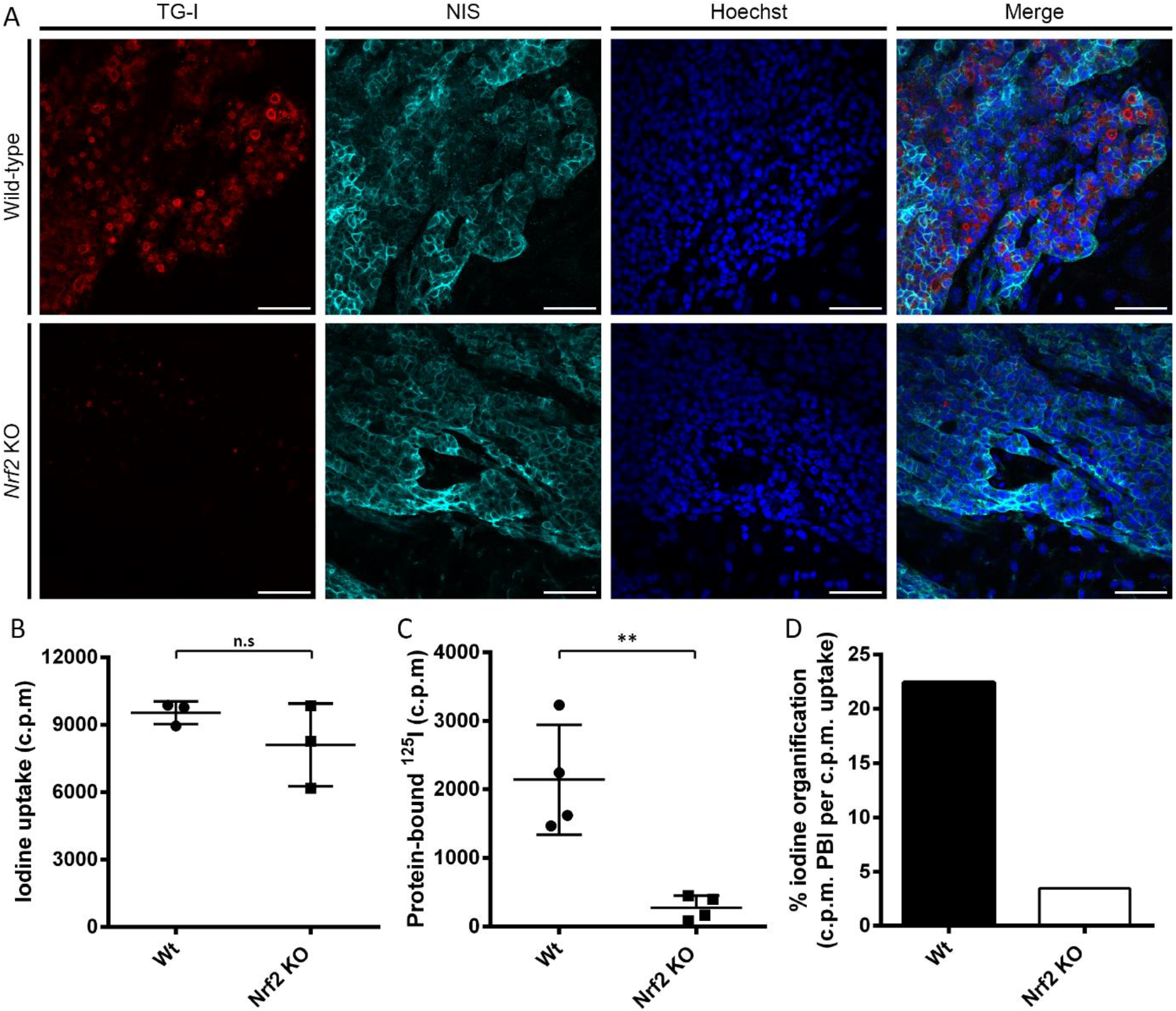
loss of Nrf2 impair thyroid function in mESC-derived thyroid follicle. Characterisation of mESC-derived thyroid organoids’ function reveals the critical need of functional Nrf2 for thyroid hormones synthesis. (A) Confocal imaging following immunofluorescence of WT and Nrf2 KO organoids after 22 days of differentiation. Samples were marked for iodinated-thyroglobulin (TG-I, Red) and NIS (Cyan). Nuclei were labelled using Hoechst (Blue). (B-D) Functional organification assay revealed equivalent ability of Nrf2 KO organoids 125I uptake (B) but reduced protein-bound 125I fraction in Nrf2 KO organoids (C), which results in a lower percentage of cells with capacity of 125I organification (D). Statistical analysis performed using Student’s t-test. Asterisks denote significant differences between conditions: **p< 0.01. “n.s”: no significant difference.

### Loss of *Nrf2* leads to important transcriptional changes in mESC-derived follicles

To better characterize the loss of the *Nrf2* gene in our mESC model and to better understand the molecular mechanisms leading to the observed phenotype, the mRNA profile of *Nrf2* WT and KO-derived thyroid follicles were compared using bulk RNA-sequencing. Importantly, although our mESC lines (*Nrf2* WT and KO) expressed the bTg-GFP reporter construct, downregulation of *Tg* expression in Nrf2 KO-derived cells prevented us from performing FACS-based cellular enrichment. Therefore, RNA sequencing experiments were performed with a mixed cell population. Initial analysis revealed significant differences between exression profiles of *Nrf2* WT and KO cells, with 1566 upregulated and 5272 downregulated genes in *Nrf2* KO condition compared with the WT (Fig. 6 A). In addition, the expression profile of key thyroid and Nrf2 pathway markers confirmed what had been previously observed by qPCR. Most of the thyroid markers were upregulated, whereas the effectors of *Tg* and Nrf2 pathway, such as *Gstp1, Gclc, Keap1, Gpx1* and *Nqo1*, were downregulated (Fig. 6B).

**Figure 6:**
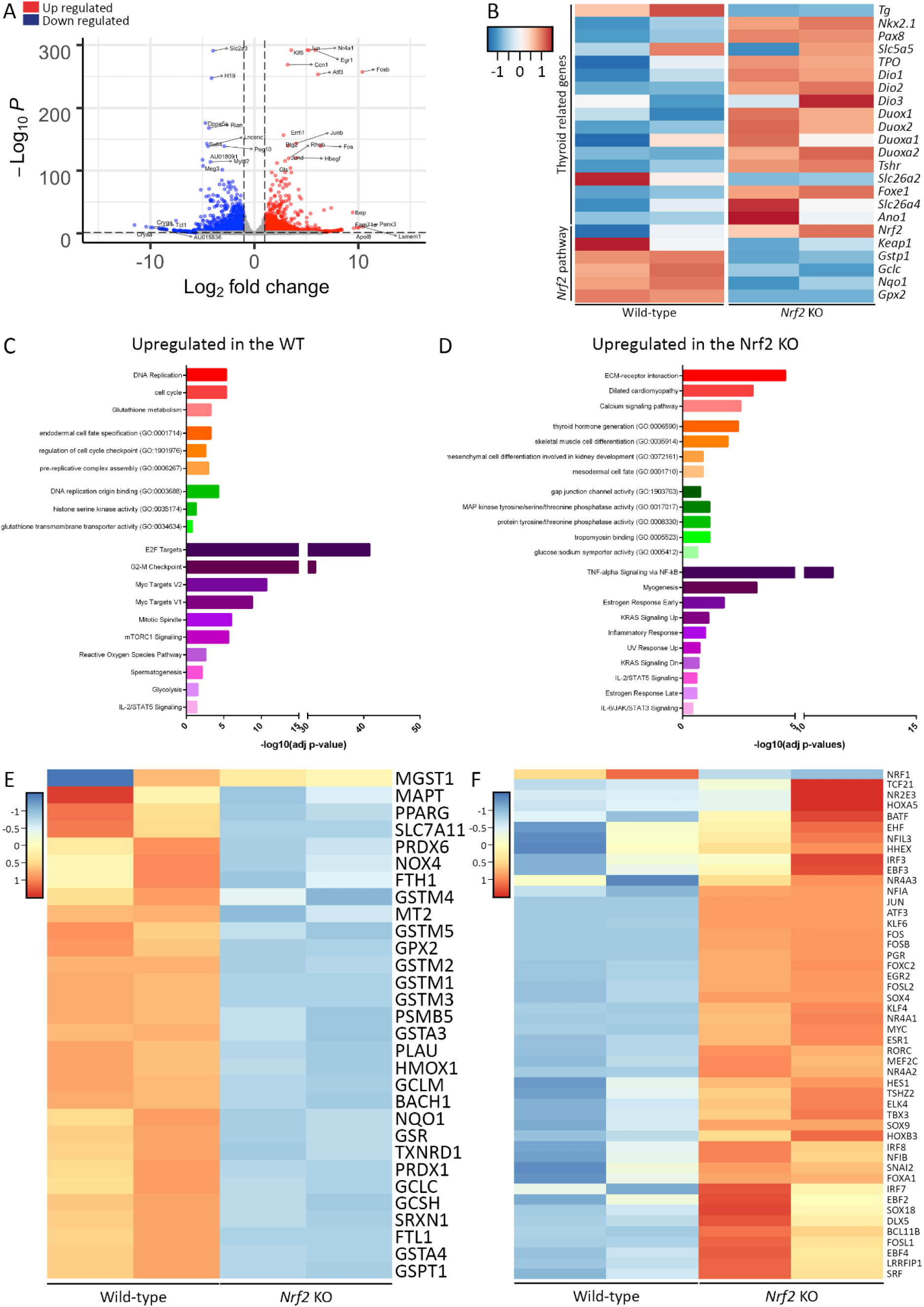
Transcriptomic profile comparison between mESC-derived Nrf2 WT and KO thyroid organoids. (A) Volcano plot showing the distribution of the down and upregulated genes among Nrf2^−/−^ cells compared to the control condition (WT). (B) Heatmap of normalized bulk RNA-Seq expression of thyroid genes and Nrf2 pathway effectors among Nrf2 WT and KO cells. Rows represent markers and columns conditions. Colour values in the heatmap represent mean expression levels. Gene ontology classification of the upregulated genes in Nrf2 WT (C) and Nrf2 KO (D) cells (Red: KEEG pathway; Orange: Biological Process; Green: Molecular Function; Violet: MSigDB Hallmark 2020). (E) Heatmap representing the expression profile of Nrf2 target genes exhibiting an antioxidant responsive element (ARE) within their promoter and/or regulatory regions. (F) Set of upregulated transcription factors among Nrf2 KO cells.

Gene Ontology (GO) classification of the most differentially expressed genes between WT and *Nrf2* KO cells, revealed that *Nrf2* expressing cells show an endodermal signature, while Nrf2 KO cells seems to express more mesodermal markers. In addition, among NRF2 WT cells, we detected upregulation of markers related to cell proliferation regulation (cell cycle regulation and DNA replication), glutathione metabolism and mTORC1 signalling. On the other hand, absence of Nrf2 results in expression signatures associated to ECM-receptor interaction, kinases activation (MAPK/RAS) and inflammation/stress (TNF-α, NF-κB, IL2 and IL6) (Fig. 6 C-D). Interestingly, these observed changes are consistent with what has been described previously in adult Nrf2 KO mice (Chartoumpekis *et al*., 2020).

As Nrf2 plays a critical role in redox homeostasis, we analysed the expression profile of *NRF2* target genes which present an antioxidant responsive element (ARE) within their promoter and/or regulatory regions (Raghunath *et al*., 2018). Validating the loss of function of *Nrf2* in our model, we demonstrate important downregulation in gene expression of this set of genes which are directly involved in several processes such as detoxification and antioxidant activity (Fig. 6E).

Aiming to identify other key genes which could be involved in the impairement of thyroid function, differential expression analysis revealed a set of transcription factors (TFs) significantly upregulated in *Nrf2* KO cells (Fig. 6F). Interestingly, identified TFs have strong interaction (Supp Fig 5) and are involved in oxidative stress response, AP-1 network and activation of several signalling pathways such as MAPK, TNF-α, NF-κB and TGF-β.

## Discussion

In this study, we evaluated the role of the *Nrf2/nrf2a* transcription factor during the development of the thyroid gland and, using zebrafish embryos and mESC-derived thyroid follicles, we highlighted its importance in both mammalian and non-mammalian thyroid development.

Using zebrafish, we identified *nrf2a* as an actor of the thyroid functional maturation using F0 Crispr/Cas9-based mutagenesis (Trubiroha *et al*., 2018). Based on these initial results we generated a stable *nrf2a* mutant line and analysed consequence of this *nrf2a* variant on thyroid development. Upon loss of *nrf2a* function, initial thyroid development occurs normally. However, functional maturation of the thyroid gland appears impaired and characterized by an enlarged non-functional thyroid gland, suggestive of a thyroid dyshormonogenesis phenotype.

Under physiological conditions, thyroid function is tightly regulated by the hypothalamo-pituitary axis via the TSH signalling pathway (Vassart and Dumont, 1992; Carvalho and Dupuy, 2017). Consistent with the lack of TH, *nrf2a* mutants showed increased levels of *tshϐ* mRNA as well as *slc5a5* and *tpo* expression. Conversely to what was previously described in zebrafish lacking TH production (Opitz *et al*., 2011; Giusti *et al*., 2020), *nrf2a* homozygous mutant embryos did not display an increased *tg* expression, instead, a slight reduction of *tg* expression could be observed. Interestingly, these results are consistent with the described control of *Tg* expression by Nrf2 in rodents (Ziros *et al*., 2018), suggesting that this role is evolutionary conserved across vertebrates.

Notably, when *nrf2a* loss of function was rescued by the tg-driven expression of a functional Nrf2a protein, we observed complete recovery of the thyroid phenotype as shown by a number of TFC and by the quantification of thyroxine production, thus demonstrating the cell-autonomous nature of the observed phenotype.

Altogether the data we obtained using zebrafish embryos show that the role of *nrf2a* goes beyond the expression control of metabolic and oxidative stress response genes and reinforce a recently published report on the role of *Nrf2* in adult mouse thyroid physiology (Ziros *et al*., 2018). In this study, P. Ziros and colleagues showed that *Nrf2* KO mice display a euthyroid state characterized by normal levels of Tsh but reduced basal levels of Tg as well as increased amount of iodinated-thyroglobulin. Although we did not analysed the thyroid gland of adult *nrf2a* KO zebrafish, this study prompted us to interrogate the role of *Nrf2* during mammalian thyroid development. Because mouse pups are developing *in utero*, we hypothesized that the phenotypic differences between *nrf2a* KO zebrafish embryos and *Nrf2* KO mice could arise from availability of maternal Tsh and T.H, thus allowing the growing pups to develop compensatory mechanisms to the loss of Nrf2 function. Therefore, in order to study the role of *Nrf2* during mammalian thyroid development without maternal compensatory effects, we decided to use our previously published mESC-derived thyroid organoids (Antonica *et al*., 2012; Romitti *et al*., 2021).

Nrf2 acts as a regulator of cellular resistance to oxidants by controlling expression of an array of antioxidant response element (AREs)-dependent genes (Buendia *et al*., 2016). As expected, we also observed that in our mouse *in vitro* model of loss-of-function, AREs genes were significantly downregulated as well as genes related to Nrf2 signalling, including Keap1 (Ziros *et al*., 2018). Importantly, this set of genes are directly involved in controlling oxidant homeostasis, detoxification and elimination of exogenous and some endogenous chemicals (Ma, 2013).

Nrf2 also acts as an anti-inflammatory molecule. Nrf2 KO mice are more likely to develop age-dependent autoimmune and inflammatory lesions (Yoh *et al*., 2001; Ma, Battelli and Hubbs, 2006). Nrf2 inhibits inflammation by inhibition of the NF-κB pathway and proinflammatory cytokine production (Li *et al*., 2008). Similar effect seems to occur in our system, since GO analysis indicated increase in TNF-α/NF-κB and inflammation signature in Nnrf2 KO condition. In addition, Nrf2 has been shown to regulate proteasomal protein degradation (Kwak *et al*., 2003), cell proliferation (Malhotra *et al*., 2010), and metabolic reprogramming (Mitsuishi *et al*., 2012), in our model exemplified by inhibition of mTORC1 signalling, cell cycle/DNA replication and glycolysis, respectively. Altogether, our results reinforce the literature data about Nrf2 role in several cellular processes as well as confirm the Nrf2 loss-of-function organoid model here presented.

Interestingly, we observed that the proportion of Nkx2.1+ cells was higher in the Nrf2 KO cells suggesting that an imbalance could occur in favour of endodermal cells within the embryonic bodies at the beginning of the differentiation protocol. This hypothesis seems unlikely considering the described role of Nrf2 in promoting mesendoderm cell fate in human and mouse embryonic stem cells (Jang *et al*., 2016; Dai *et al*., 2020), which was here confirmed by GO analysis of our transcriptomic data indicating mesodermal signature in our Nrf2 KO RNA sequencing analysis. Another hypothesis could be that cells committed toward thyroid fate undergo higher proliferation rate when lacking functional Nrf2. Nrf2 is well known to regulate the quiescence, proliferation and differentiation rate of various stem and progenitor cell type (Dai *et al*., 2020). This role is best exemplified in haematopoietic stem and progenitor cells (HSPCs) which demonstrated that Nrf2 was a critical factor to maintain the stemness and the quiescence (Tsai *et al*., 2013). Recently, Murakami and colleagues demonstrated that an over-activation of the Nrf2 signalling was as detrimental to the stemness, quiescence and longevity of the HSPCs as the lack of functional Nrf2 (Murakami *et al*., 2017). The authors propose that the role of Nrf2 in maintaining the HSPCs depends on a fine regulation of the Nrf2 activity, over-expression or lack of Nrf2 leading to a higher proliferative and differentiation rate ultimately causing the exhaustion of the HSPC niche (Murakami *et al*., 2017). On the other hand, RNA sequencing indicated a higher proliferative signature in Nrf2 WT cells, which here could be due to the higher proportion of non-thyroid cells in that condition. Thus, the mechanism inducing this higher proportion of Nkx2.1+ cells remains unknown and further studies will be necessary to clarify it.

Nrf2 KO mESC-derived thyroid cells showed no defect in the uptake of iodine but were unable to organify it and produce iodinated-thyroglobulin, similar to what we observed in *nrf2aΔ5* mutant zebrafish embryos, which may be due to the lack of *Tg* production.

The most striking difference we observed between the two models was the absence of follicular organisation in *Nrf2* KO thyroid cells. Thyroid folliculogenesis is a poorly understood process and recent studies suggest that thyroid follicle formation occurs in two steps. The first step is endothelial cell-dependent basal laminal formation around unpolarised thyroid cells (Villacorte *et al*., 2016; Pierreux, 2021). The second step is the formation of microlumen through the migration of thyroglobulin-filled microvesicles to the opposite side of the basal lamina (Gonay *et al*., 2021). Although follicular lumen expansion is a thyroglobulin-independent process (Liang *et al*., 2018), it is possible that the initial microlumen formation is thyroglobulin-dependent, although more studies are necessary to confirm it.

In zebrafish, only few studies have described thyroid folliculogenesis and showed that soon after budding out of the endoderm, thyroid cells are already organised as a multicellular follicle-like structure (Alt *et al*., 2006; Porazzi *et al*., 2009; Opitz *et al*., 2012). In addition, zebrafish thyroid gland is able to produce thyroid hormones while proliferating and migrating toward its final location which greatly differs from mammalian thyroid development (Opitz *et al*., 2011, 2012; Nilsson and Fagman, 2017).

Evolutionarily, this difference is thought to be caused by the earlier need for thyroid hormones in zebrafish embryos compared to the mouse embryos. Indeed, as soon as it hatches (between 2 and 3 days of age) the zebrafish embryos starts to feed on prey, which requires the maturation of its gastro-intestinal apparatus, the inflation of its swim bladder and the maturation of its craniofacial cartilages. All these processes are controlled by thyroid hormones (Liu and Chan, 2002; Bagci *et al*., 2015; Campinho, 2019). In this context, although it is currently admitted, but not proven, that zebrafish and mouse folliculogenesis are following similar mechanisms, our data might suggest that this process could be Tg-dependent in mESC-derived thyroid organoids and Tg-independent in zebrafish thyroid development.

Another difference between the thyroid phenotype of the two models concerns the extent of thyroglobulin expression. Although both models show dysregulation of thyroglobulin expression, the Nrf2 KO thyroid cells showed very low *Tg* expression whereas the *nrf2aΔ5* mutant zebrafish showed only a slight reduction in *tg* expression. One possible explanation could be the availability of Tsh as a signalling molecule. Conversely to our zebrafish model, where thyroid function is regulated by the hypothalamo-pituitary-thyroid axis, mESC-derived thyroid organoids are not stimulated with Tsh during the differentiation protocol. Instead, we used cAMP (Romitti *et al*., 2021) as an alternative to Tsh as it is known to mediate most of Tsh signalling effects in thyroid cells, from hormone biosynthesis and release to thyrocyte proliferation and differentiation (Ortiga-Carvalho *et al*., 2016). The role of Tsh signalling in controlling thyroglobulin expression is still a matter of debate. While some studies have demonstrated the TSH-independent onset of thyroglobulin expression in mouse (Marians *et al*., 2002; Postiglione *et al*., 2002; Christophe, 2004), others have suggested that TSH signalling could play a role in the maintenance of Tg expression, in adult rats and dog thyroids (van Heuverswyn *et al*., 1984; Gérard, Roger and Dumont, 1989; Christophe, 2004). In zebrafish embryos, inhibition of Tsh signalling through *tshr* knock-down induced a reduction of thyroglobulin expression (Opitz *et al*., 2011). Although this reduction is not as striking as what was observed for *slc5a5* and *tpo* expression in *tshr* morphants, it suggests that, despite not being not completely indispensable, Tsh signalling modulates thyroglobulin expression in zebrafish.

At the transcriptional levels, analysis revealed a set of TFs upregulated in Nrf2 KO cells. Among those TFs, several are important effectors of AP-1 network, such as *Atf, Jun* and *Fos* family. AP-1 transcription factors play an important role in controlling of cell proliferation, survival and death (Karin, Liu and Zandi, 1997) and several mechanisms account for its stimulation such as, growth factors, proinflammatory cytokines and UV radiation (Karin, 1995). It is suggested that most important mediator of the growth factor response is likely to be the ERK/MAP kinase (MAPK) cascade while the responses to proinflammatory cytokines and UV radiation are mostly dependent on two other MAPK cascades, JNK and p38 (Karin, 1995). Interestingly, we also observed increase of MAPK expression signature in our model, which might suggest that the effect of Nrf2 loss on thyroid function might be promoted by hyperactivation of AP-1 network, in response to the changes in the oxidative cell state.

Interestingly, we also observed a difference in the expression of our Tg reporter systems upon loss of *Nrf2/nrf2a* function. In these systems, Tg expression is reported either by EGFP driven by the bovine Tg promoter in mESC-derived organoids (Antonica *et al*., 2012; Romitti *et al*., 2021) or by EGFPnls driven by the zebrafish *tg* promoter in zebrafish embryos (Trubiroha *et al*., 2018). However, while the loss of Nrf2 led to a drastic reduction of EGFP expression in mESC-derived organoids, the loss of *nrf2a* in zebrafish embryos did not affect EGFPnls expression in thyroid cells. As it was previously shown that rodent and humanTg is directly controlled by Nrf2 through two evolutionarily conserved AREs located in its distal enhancer (Ziros *et al*., 2018), we examined whether these ARE are conserved in the bovine and zebrafish Tg regulatory sequences. Whereas the two AREs are present in the bovine Tg regulatory sequence, they are absent from the zebrafish one (Supp Fig. 6). This suggests that direct control of Tg by the binding of Nrf2 in the distal Tg enhancer is evolutionarily conserved in mammalian vertebrates but not in non-mammalian vertebrates and indicates that the observed discrepancy between the two reporter systems is due to the absence of AREs in the zebrafish *tg* promoter.

In summary, our data demonstrate for the first time the evolutionarily conserved importance of nrf2/nrf2a in thyroid development in mammals and other vertebrates. Using a combination of in vivo and in vitro models, the present study characterized the effect of nrf2 loss of function on thyroid organogenesis at the morphological and transcriptomic levels. While our study highlighted the lack of thyroglobulin expression as the main cause of the observed phenotypes, it also suggested that this is driven by species-specific mechanisms. While direct control of Tg expression by nrf2 has now been clearly demonstrated in mice, our data suggest that this direct control appears to be absent in zebrafish. Further studies will be necessary to unravel the mechanism by which nrf2a acts on thyroglobulin expression in zebrafish. In addition, it would be interesting to analyze the role of nrf2 during thyroid organogenesis in a broader range of models, thereby expanding our knowledge of the molecular mechanisms that control thyroid development in vertebrates.

## Materials and methods

### Animal husbandry

Zebrafish embryos were collected by natural spawning of adult individuals, embryos and larvae were raised at 28.5 °C under standard conditions (Westerfield, 2000) and staged according to hours (hpf) or days post fertilization (dpf) as described (Kimmel *et al*., 1995). Embryos were collected and raised in 90mm petri dishes containing 25 mL of embryo medium. At 1 dpf, all embryos were dechorionated by a 20 min treatment with 0.6 mg/mL pronase (Sigma). Screening and live imaging of embryos and larvae were performed after anaesthesia using 0.02% tricain (Sigma) The following zebrafish lines were used in this study: wild-type AB strain (WT), Tg(*tg:nlsEGFP*)^ulb4^ (Trubiroha *et al*., 2018), *nrf2aΔ5* mutant generated by Crispr/Cas9-based mutagenesis (this study) and Tg(*tg:nrf2a_T2A_mKO2nls*) (this study). All zebrafish husbandry and all experiments were performed under standard conditions in accordance with institutional (Université Libre de Bruxelles-ULB) and national ethical and animal welfare guidelines and regulation. All experimental procedures were approved by the ethical committee for animal welfare (CEBEA) from the Université Libre de Bruxelles (protocol 578 N).

### Single-guide RNA design and synthesis

Design and generation of single guide RNA (sgRNA) used in this study were performed as previously described (Trubiroha *et al*., 2018). Briefly, The Sequence Scan for CRISPR software (http://crispr.dfci.harvard.edu/SSC/) was used to identify sgRNA sequences with predicted high on-target activity (Xu *et al*., 2015). DNA templates for sgRNA synthesis were produced using the PCR-based short-oligo method described by Talbot and colleagues in 2014 (Talbot and Amacher, 2014). DNA templates for *in* vitro transcription of sgRNA were generated by annealing a scaffold oligonucleotide (sequence: AAAGCACCGACTCGGTGCCACTTTTTCAAGTTGATAACGGACTAGCCTTATTTTAACTTGCTATTTCTAGCTCTA AAAC) with a gene-specific oligonucleotides containing the SP6 promoter sequence (GCGATTTAGGTGACACTATA), a 20 base target sequence (see Supplementary Table 1) and a complementary sequence to the scaffold oligo (GTTTTAGAGCTAGAAATAG). PCR amplification was performed using Taq PCR Core Kit (QIAGEN). Reaction mix contained 20 nM scaffold oligo, 20 nM gene-specific oligo, and 260 nM of each universal flanking primer (forward: GCGATTTAGGTGACACTATA, reverse: AAAGCACCGACTCGGTGCCAC). Purification of the PCR products were performed using MinElute Reaction Cleanup kit (QIAGEN) followed by a phenol-chloroform-isoamylalcohol extraction. sgRNAs were synthesized in vitro from purified PCR products by using the SP6 RNA-polymerase (NEB). 20 μL reactions contained 1 μg DNA template, 1x reaction buffer, 2.5 mM of each rNTP (Roche), 40 U RNase Inhibitor (Thermo Fisher Scientific), and 6 U Large Fragment of DNA Polymerase I (Thermo Fisher Scientific). After pre-incubation for 15 min at room temperature, 20 U SP6 RNA-polymerase were added and reactions were run over night at 40 °C. After treatment with 3 U DNase I (AmpGrade; Thermo Fisher Scientific), sgRNAs were purified using High Pure PCR Cleanup Micro Kit (Roche). Concentration of sgRNAs was measured by Nanodrop (Thermo Fisher Scientific) and integrity was checked by gel electrophoresis. Aliquots of sgRNA solutions were stored at −80 °C until use.

### sgRNA and Cas9 injection

Injection mix containing Cas9 protein (PNA Bio; 100 ng/μl final concentration), sgRNA (80 ng/μL final concentration), 200 mM KCl and up to 0.15% phenol red (Sigma) was injected into one-cell stage embryos. Approximately 3 nL of active sgRNA-Cas9 ribonucleoprotein complex were injected in each one-cell stage Tg(tg:nlsEGFP) embryo. For each sgRNA, at least three independent injection experiments were performed with spawns from at least three different founder fish.

### Phenotypic analysis of F0 crispants and stable mutant embryos

Following anaesthesia using 0.02% tricaine, thyroid development was monitored in live zebrafish at 55 hpf, 80 hpf and 6 dpf by visual inspection using a M165 FC fluorescence stereomicroscope (Leica). For F0 crispants analysis, all injected and non-injected embryos were inspected for gross developmental defects, deviations in size, shape or location of the thyroid, and for the overall intensity of the fluorescent reporter signal. For imaging of the thyroid phenotypes, embryos and larvae were immobilized in 1% low-melting point agarose (Lonza) containing 0.016% tricaine and positioned in Fluoro-Dish glass bottom dishes (WorldPrecisition Instruments). Imaging was performed using a DMI600B epifluorescence microscope equipped with a DFC365FX camera and LAS AF Lite software (Leica). If indicated, confocal live imaging was performed with a LSM780 confocal microscope (Zeiss) using Zen 2010 D software (Zeiss).

### Genotyping of zebrafish *nrf2aΔ5* mutants

Genomic DNA was extracted by a 10min lysis treatment with 50mM NaOH at 95°C either from adult tailfin clips or from whole larvae following. Subsequently, lysate was neutralized using 0.5M TRIS (Ph 8.0) and used as template PCRs using primers spanning the mutated site. The following primer sequences were used: Forward 5’-TGATCATCAATCTGCCCGTA-3’ and Reverse 5’-TCTCTTTCAGGTTGCTGCTG-3’. PCR product were then analysed by Sanger sequencing to identify wild-type, heterozygous and homozygous carriers.

### Phenylthiourea treatment of zebrafish embryo

Zebrafish embryos and larvae were treated with phenylthiourea (PTU; Sigma) from 24 hpf onwards in order to induce hypothyroid conditions (Elsalini and Rohr, 2003; Opitz *et al*., 2011). A stock solutions of 100 mg/L PTU was diluted in embryo medium to make experimental treatment solutions of 30 mg/L PTU. A control group was maintained in embryo medium. Embryos and larvae were treated in 90 mm petri dishes containing 25 mL of treatment solution and solutions were renewed every other day.

### Whole-mount immunofluorescence (WIF)

144 hpf larvae were fixed in 4% PFA (pH 7.3) over night at 4 °C and stored in PBS containing 0.1% Tween-20 at 4 °C until use. WIF experiments were performed as previously described (Opitz *et al*., 2011) using the following antibodies: Goat anti-T4 polyclonal antibody (1:1000; Byorbyut), chicken anti-GFP polyclonal antibody (1:1000; Abcam), Mouse anti-iodinated thyroglobulin polyclonal antibody (1:1000; Dako), cy3-conjugated donkey anti-goat IgG antibody (1:250; Jackson ImmunoResearch), Alexa Fluor 488-conjugated goat anti-chicken IgG antibody (1:250; Invitrogen) and cy5-conjugated donkey anti-mouse IgG antibody (1:250, Jackson ImmunoResearch). Following immunostaining, samples were postfixed in 4% PFA and gradually transferred to 100% glycerol for conservation at 4°C. Phenotypical analysis was performed using a M165 FC fluorescence stereomicroscope (Leica). Whole mount imaging of stained specimens was performed either with a DMI600B epifluorescence microscope (Leica) equipped with a DFC365FX camera or, if stated, using a LSM780 confocal microscope (Zeiss).

### Quantitative analysis of T4 and Tg-I fluorescent signal intensities

Quantification of fluorescent signal was performed as previously described (Thienpont *et al*., 2011), whole mount imaging of the thyroid region was performed on stained samples using a DMI600B epifluorescence microscope. Using LAS AF imaging software (Leica), pixel sum values were obtained for each thyroid follicular lumen (region of interest 1 to X, ROI 1-X) and adjacent non-fluorescent regions (ROI-bg, background). For each embryo, the intensity of each ROI 1-X was corrected by subtracting the intensity of ROI-bg. The sum of each individual background-corrected fluorescence signal intensities of a single embryo constitutes the measurement of either its T4 or Tg-I content. Values from individual fish were used in comparisons between treatment or mutant groups and controls.

### Generation of the IscE1 Meganuclease-based Tg(*tg:nrf2a_T2A_mKO2nls*) transgenic zebrafish line

The Gateway cloning technology-based Tol2kit (Kwan *et al*., 2007) was used to generate a rescue construct in which the functional *nrf2a* coding sequence fused to a nuclear localised mKO2 fluorescent protein with a viral T2A linker peptide (Kim *et al*., 2011) is expressed under the control of a fragment of the zebrafish *tg* gene promotor previously shown to allow thyroid-specific transgene expression (Opitz *et al*., 2012). Generation of the plasmid was essentially performed as previously described (Trubiroha *et al*., 2018). The full nrf2a coding sequence fused to the mKO2 protein via the T2A peptide was synthesized by Genewiz company qnd was subsequently cloned into a pME vector to create a pME-nrf2a_T2A_mKO2nls vector. Assembly of the final Tol2Tg(*tg:nrf2a_T2A_mKO2nls*) vector was performed using a LR reaction (Villefranc, Amigo and Lawson, 2007).

Capped mRNA encoding for transposase was generated by in vitro transcription of NotI-linearized pCS-zT2TP plasmid (Kawakami, 2004) using mMessage mMachine SP6 kit (Ambion). Transgenic embryos were generated by the co-injection of Tol2Tg(*tg:nrf2a_T2A_mKO2nls*) vector (25 ng/μL) and Tol2 transposase mRNA (35 ng/μL) in one-cell stage *nrf2aΔ5* heterozygous embryos. Injected embryos were screened at 55 and 80 hpf for mKO2 expression in thyroid cells and only F0 fish presenting thyroid-specific mKO2 expression were raised to adulthood to identify potential germline transmitter. F0 founders with germline transmission were out-crossed with WT fish to generate stable F1 transgenic Tg(*tg:nrf2a_T2A_mKO2nls*) animals. Transgenic F1 embryos were raised to adulthood and genotyped for the presence of the *nrf2aΔ5* mutation. Tg(*tg:nrf2a_T2A_mKO2nls*) line was maintained in a WT and in a *nrf2aΔ5* mutant genetic background.

### Generation of the A2lox_Nkx2-1_Pax8_Nrf2^−/−^ mouse ESC line using CRISPR/Cas9

SgRNAs targeting mouse Nrf2 coding sequence were selected using the prediction algorism CRISPR, as described above. In order to compare the effect of Nrf2 knockout in mESC-derived thyroid with zebrafish nrf2aΔ5 mutants, the selected gRNA targets the same genomic region chosen for zebrafish, the DNA binding and transactivation of Nrf2 gene. The most suitable sgRNA was chosen and checked for possible off-targets. Initially, gRNA sequence (GTGTGAGATGAGCCTCTAAG) was cloned inside the pU-BbsI-T2A-Cas9-BFP plasmid (64323, Addgene). Then the A2Lox.Cre_TRE-Nkx2-1/Pax8_Tg-EGFP mouse ESCs (Romitti, 2021) were transfected using Lipofectamine 3000 reagent (Invitrogen), and after 24 h, BFP-expressing cells were sorted, clonal seeded, expanded and analysed by PCR and sequenced. For that, genomic DNA was extracted from individual mESCs clones using the DNeasy Blood and Tissue Kit from Qiagen (Qiagen). PCR was performed using 100 ng of genomic DNA and Q5 High-Fidelity Taq Polymerase (New England Biolabs) was used according to the manufacturer’s instructions. Primer sets (Fw: 5’-TCCTATGCGTGAATCCCAAT-3’ and Rv: 5’-TAAGTGGCCCAAGTCTTGCT-3’) were used to amplify the Crispr/cas9 target region. PCR products were cloned into TOPO-Blunt plasmid by using Zero Blunt^®^ PCR Cloning Kit (Thermo Fisher) and sequenced using Sanger method. Clones were chosen based on the identification of frameshift mutations in *Nrf2* gene. For thyroid differentiation, two different clones were selected and validated according to the maintenance of pluripotency, tested by spontaneous differentiation into the three germ layers and induction of Nkx2-1-Pax8 transgenes under Dox stimulation.

### mESC_Nkx2-1_Pax8_Nrf2^−/−^ culture and thyroid differentiation

Modified mESCs were_cultured and differentiated as previously described (Antonica *et al*., 2012, 2017). Briefly, cells were cultured on irradiated mouse embryonic fibroblasts feeder-layer using a mESC medium (Antonica et al 2012, 2017). For thyroid differentiation induction, Nkx2-1_Pax8_Nrf2^−/−^ and Nkx2-1_Pax8_Nrf2^wt/wt^ (control) mES cells were cultured for four days in suspension using hanging drops method in order to generate embryoid bodies (EBs). After four days, generated EBs were collected and embedded in 50μl (around 40 EBs) of growth-factor-reduced Matrigel® (BD Biosciences), plated in 12-well plates and treated for three days with 1 mg/ml Doxycycline (Sigma), in order to stimulate Nkx2-1 and Pax8 transgenes expression. After Dox withdraw, cells were treated for two weeks with 300μM 8-br-cAMP (Biolog Inc.) or 1mU/ml rhTSH (Genzyme) (Romitti *et al*., 2021). Samples were collected at distinct time points for RNA expression and histological analysis.

### Quantitative PCR (qPCR) Analysis

qPCR was performed on cDNA generated from thyroid organoids derived from *Nrf2* WT and KO cells at differentiation day 7 and 22 or from whole zebrafish embryos at 6 days of age. Total RNA was extracted from samples using RNeasy micro kit (Qiagen) according to the manufacturer’s instructions. cDNA was generated by reverse transcription using Superscript II kit (Invitrogen). qPCR was performed in triplicates using Takyon (Eurogentec) and CFX Connect Real-Time System (Biorad).

Results are presented as linearized values normalized to the housekeeping gene, B2microglobulin and the indicated reference value (2-DDCt). The gene expression profile was obtained from at least three independent samples and compared to their respective control (Nkx2-1_Pax8_Nrf2^wt/wt^ mESC line or WT zebrafish embryos). Primers sequences are listed in the table 1.

**Table 1:**
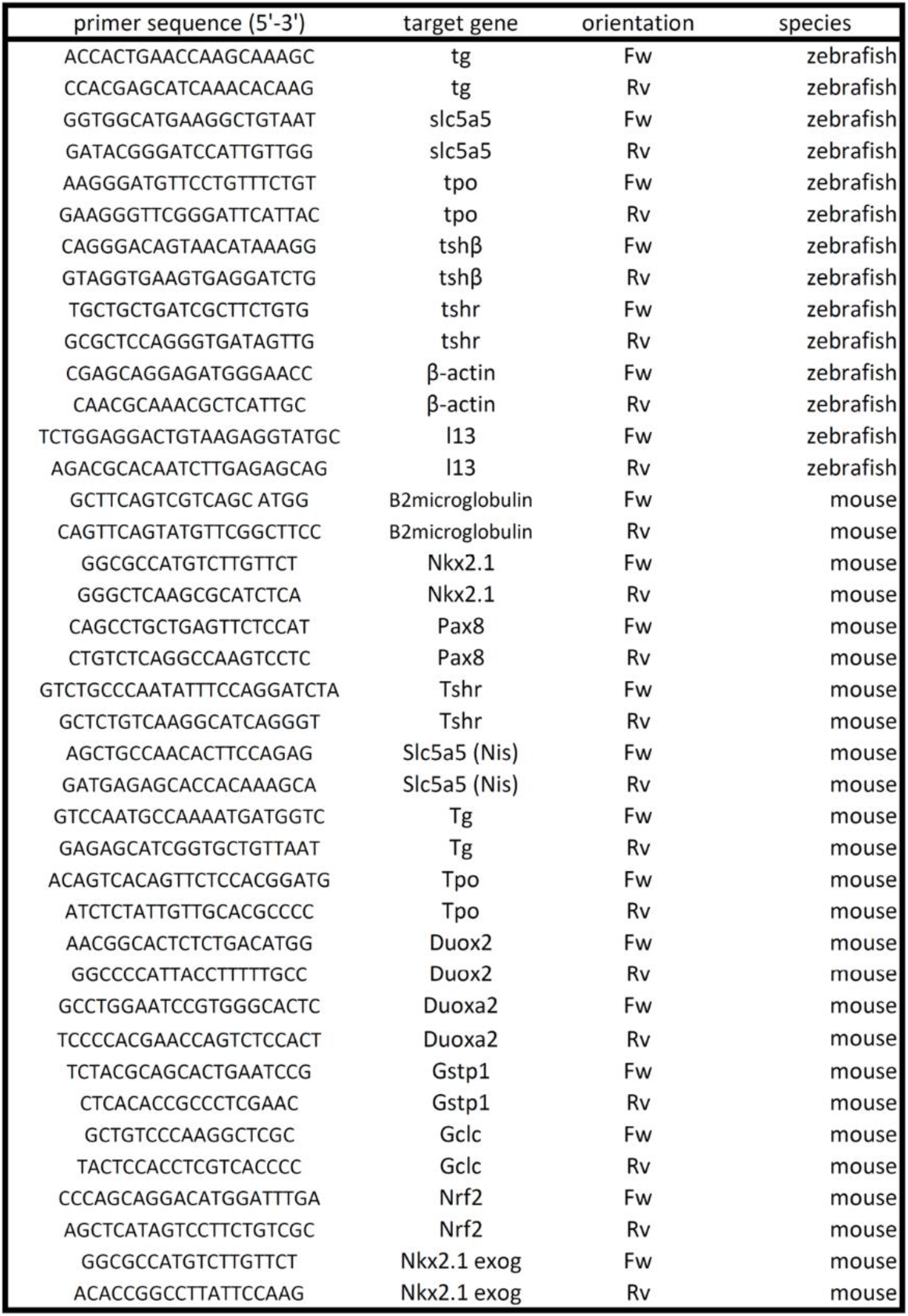
qPCR primer table. List of the primers used to perform qPCR analysis.

### Flow cytometry intracellular immunostaining

Nrf2^wt/wt^ and Nrf2^−/−^ - derived cells were collected at day 22 (cAMP condition) and prepared for flow cytometry immunostaining as follows: Matrigel-embedded organoids were digested using a HBSS solution containing 10 U/ml dispase II (Roche) and 125 U/ml of collagenase type IV (Gibco, Thermo Fisher) for 30-60 minutes at 37°C; then, enzymes were inactivated using 10% FBS followed by incubation with TripLE Express solution (Thermo Fisher) for 10-15 min, at 37°C, to dissociate remaining structures (including thyroid follicles) and finally obtain single cell suspension. After inactivation by addition of differentiation medium, cells were centrifuged, rinsed with PBS and fixed in 1.6% PFA solution in PBS for 15min at RT, followed by cell permeabilization with 0.1% Triton solution in PBS for 15 min at 4°C under agitation. After centrifugation, cells were blocked using 4% horse serum (HS) and 0.5% Tween 20 PBS blocking solution for 10 min (4°C under agitation). Primary anti-rabbit Nkx2-1 antibody (1:100) was diluted in blocking solution and samples incubated for 30 min (4°C under agitation). Cells were then rinsed three times with washing solution (0.5% BSA and 0.5% Tween in PBS) and then incubated with Cy5-conjugated anti-rabbit antibody (1:300, in blocking solution0 for 30 min (4°C under agitation). Data concerning bTg-GFP (endogenous expression) and Nkx2-1 expression were obtained and processed using a LSRFortessa X-20 flow cytometer and FACSDiva software (BD Biosciences), respectively. Unstained cells and isotype controls were included in all experiments. In addition, GFP+ cell quantification was used to estimate the proportion of Tg+ cells generated by Nrf2^wt/wt^ and Nrf2^−/−^ cells.

### Immunofluorescence

Protein immuno-detection experiments were performed as previously described (Antonica *et al*., 2012; Romitti *et al*., 2021). Briefly, organoids were fixed in 4% paraformaldehyde (Sigma) for 1 h at room temperature (RT), washed three times in PBS and blocked in a solution containing 3% bovine serum albumin (BSA; Sigma), 5% horse serum (Invitrogen) and 0.3% Triton X-100 in PBS (Sigma) for 30 min at RT. The primary and secondary antibodies were diluted in a PBS solution containing 3% BSA, 1% horse serum and 0.1% Triton X-100. Primary and secondary antibodies dilutions and specifications are described at table 2 and table 3 respectively. Coverslips containing the stained organoids were mounted with Glycergel (Dako) and samples were imaged on Leica DMI6000 or Zeiss LSM 780 confocal microscope.

### Iodide Organification Assay

Nrf2^wt/wt^ and Nrf2^−/−^ - derived organoids (cAMP treated) were tested for the ability of Iodide uptake and organification at differentiation day 22. Matrigel drops containing approximately same number of organoids were initially washed with HBSS and incubated with 1 ml of a HBSS solution containing ^125^I (1,000,000 c.p.m./ml; PerkinElmer) and 100 nM of sodium iodide (NaI; Sigma) for 2 h at 37°C. After incubation, 1 ml of 4 mM methimazole (MMI), was added and cells washed with ice-cold PBS. In order to collect and dissociate the organoids from the MTG drops, cells were incubated with 0.1% trypsin (Invitrogen) and 1 mM EDTA (Invitrogen) in PBS for 15 min. For iodide uptake evaluation, cells were collected in polyester tubes and radioactivity was measured with a c-counter. After, proteins were precipitated using 1 mg of gamma-globulins (Sigma) and 2 ml 20% TCA followed by centrifugation at 2,000 r.p.m. for 10 min, at 4°C and the ^125^I protein-bound (PBI) was measured. Iodide organification was calculated by iodide uptake/PBI ratio and the values expressed as a percentage. As control for iodide uptake and protein-binding cells were also treated with 30mM sodium perchlorate (Nis inhibitor; NaClO4, Sigma-Aldrich)) and 2 mM methimazole (TPO inhibitor; MMI, Sigma-Aldrich), respectively. Experiments were performed using at least three independent replicates.

### Bulk RNA-Sequencing

Bulk RNA-seq was performed in Nrf2^wt/wt^ and Nrf2^−/−^-differentiated cells at day 22 of our differentiation protocol. Total RNA isolation was performed as described above (qPCR section). The RNA concentration and quality was accessed using Bioanalyser 2100 (Agilent) and RNA 6000 Nano Kit (Agilent). RNA integrity was preserved, and no genomic DNA contamination was detected. Illumina TruSeq RNA Library Prep Kit v2 was employed, as indicated by the manufacturer, resulting in high quality indexed cDNA libraries, which were quantified using Quant-iT PicoGreen kit (Life Sciences) and Infinite F200 Pro plate reader (Tecan); DNA fragment size distribution was examined on 2100 Bioanalyser (Agilent) using DNA 1000 kit (Agilent). The multiplexed libraries (10ρM) were loaded on flow cells and sequenced on the HiSeq 1500 system (Illumina) in a high output mode using HiSeq Cluster kit v4 (Illumina). Roughly 10 million paired-end reads were obtained per sample. After removal of low-quality bases and Illumina adapter sequences using Trimmomatic software (Bolger, Lohse and Usadel, 2014), sequence reads were aligned against mouse reference genome (Grcm38/mm10) using STAR software with default parameters (Dobin *et al*., 2013). Raw counts were obtained using HTSeq software (Anders, Pyl and Huber, 2015) using Ensembl genome annotation. Normalization, differential expression and Gene Ontology analyses were performed using two biological replicates per sample, using website iDEP version 0.93 (Ge, Son and Yao, 2018). Genes in which the expression levels were lower than 5 were filtered out.

### Regulatory sequence analysis

The regulatory sequence for mouse Tg was downloaded from the EPDnew database (http://epd.vital-it.ch, Promoter ID: TG_1) (Dreos *et al*., 2017). Bovine and zebrafish Tg regulatory sequences used in this study are the same as the one used to develop the previously published reporter mESC and zebrafish lines (Antonica *et al*., 2012; Trubiroha *et al*., 2018). Identification of potential ARE sequences was done through the JASPAR database (http://jaspar.genereg.net) (Mathelier *et al*., 2016) using the ARE weight matrix MA0150.2. Only identified sequences with a p-value and a q-value <0.05 were considered for the analysis.

## Acknowledgment

We thank members of the Costagliola laboratory for comments on the manuscript and technical assistance and members of IRIBHM fish facility for technical assistance. We thank J.-M. Vanderwinden from the Light Microscopy Facility and Christine Dubois from the FACS facility for technical assistance at ULB. Bulk RNA sequencing was performed at the Brussels Interuniversity Genomics High Throughput core (BRIGHTcore: www.brightcore.be).

---

**Supplementary figure 1:**
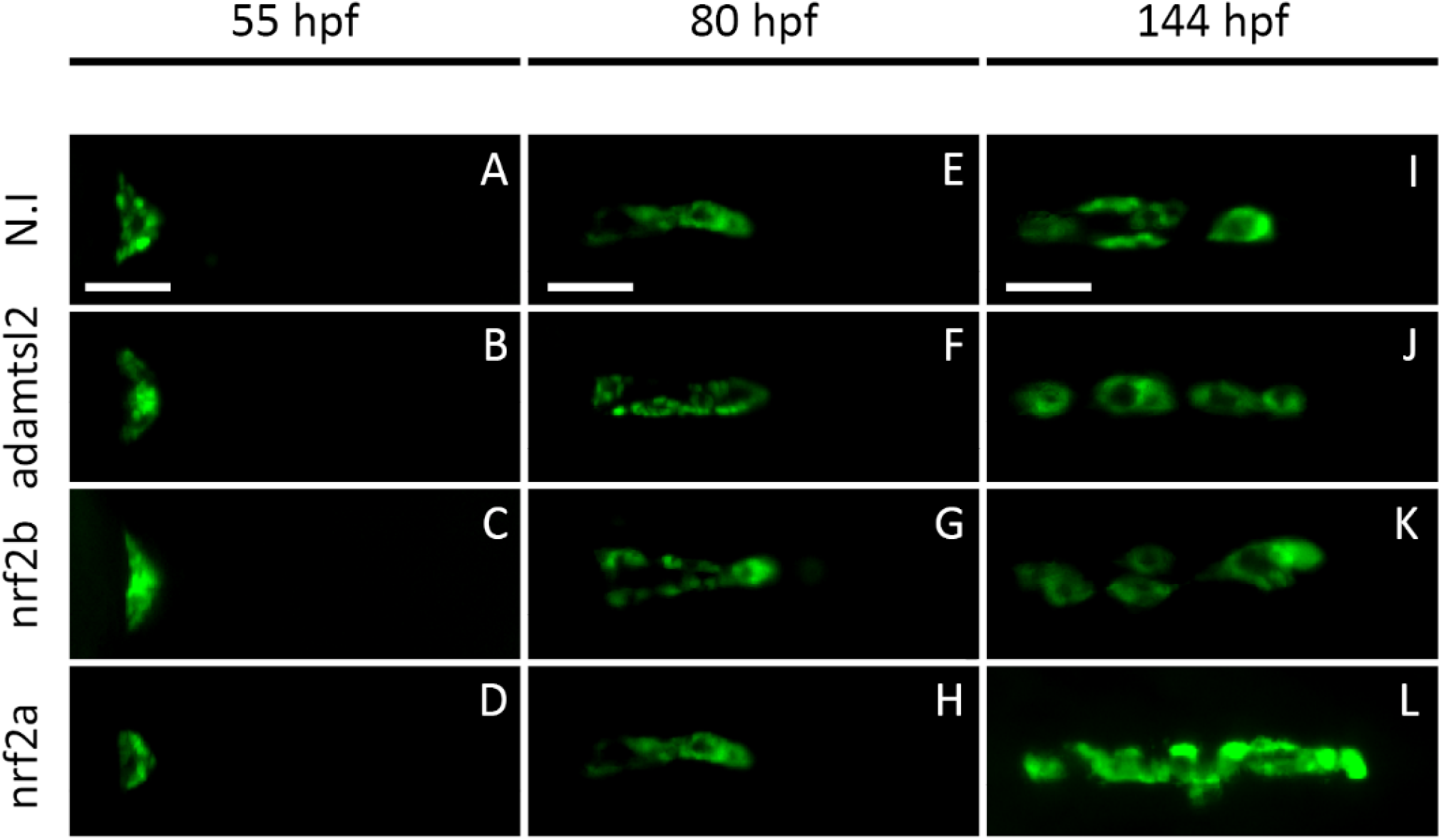
F0 crispants screening approach identifies *nrf2a* as an actor of late thyroid development in zebrafish. Live imaging of transgenic Tg(tg:nlsEGFP) zebrafish embryos from 55 to 144 hpf following one-cell stage injection of single-guide RNA and Cas9 protein targeting either *nrf2a, nrf2b* or *admatsl2*. Epifluorescence live imaging showed that early stages of thyroid development are not affected in *nrf2a* and *nrf2b* crispants embryos when compared to their *adamtsl2* crispants and WT siblings. However, at 6dpf *nrf2a* crispants displayed enlargement of their thyroid gland compared to their *nrf2b* and *adamtsl2* crispants and non-injected siblings. (A-L) show pictures of the thyroid region of a representative embryo at specified developmental stage (ventral view, anterior is to the right). Scale bars: 100μm.

**Supplementary Figure 2:**
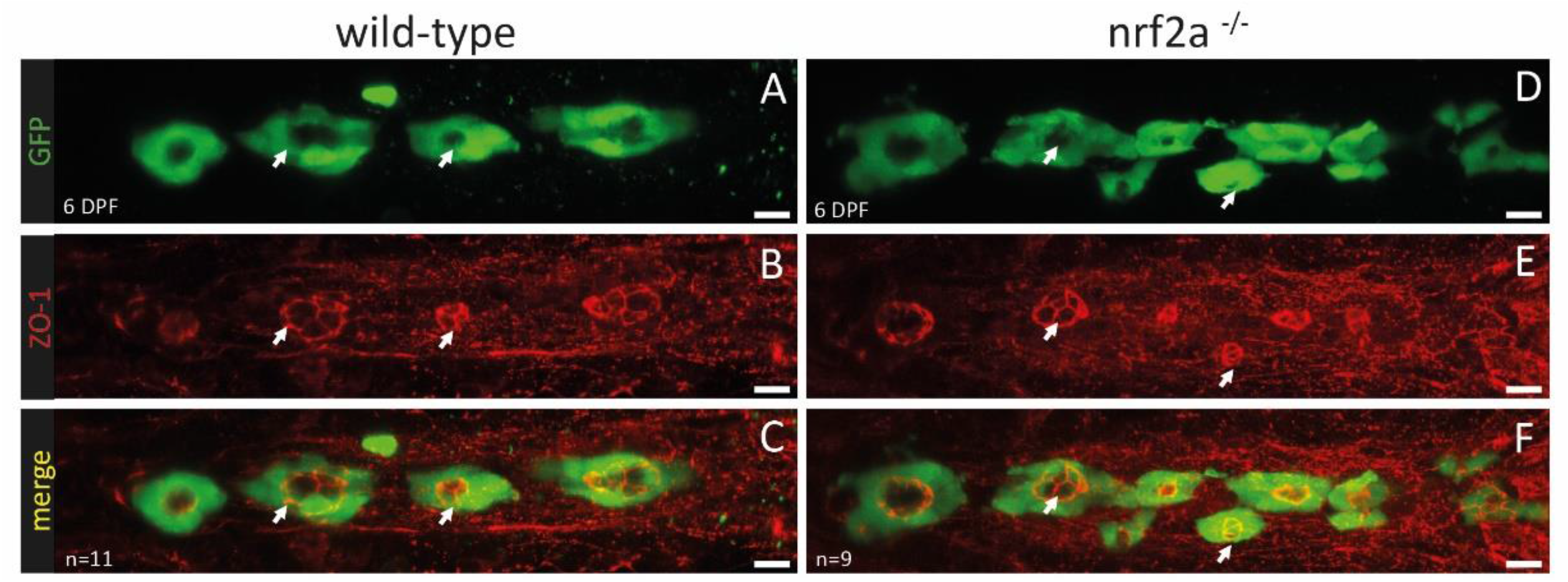
Loss-of-function of *nrf2a* does not impair thyroid follicles polarisation. Immunofluorescence on 6dpf Tg(*tg:nlsEGFP*) zebrafish embryos using antibodies targeting EGFP and ZO-1 (Cy3 - Red) reveals normal follicular polarisation in *nrf2a* homozygous mutant compared to their wild-type siblings (A-F). Upper row shows GFP staining only, middle row shows ZO-1 staining only and last row shows a merge of both staining. White arrows are showing representative follicles in both wild-type and homozygous mutant. Scale bars 10μm. Ventral view, anterior to the left.

**Supplementary Figure 3:**
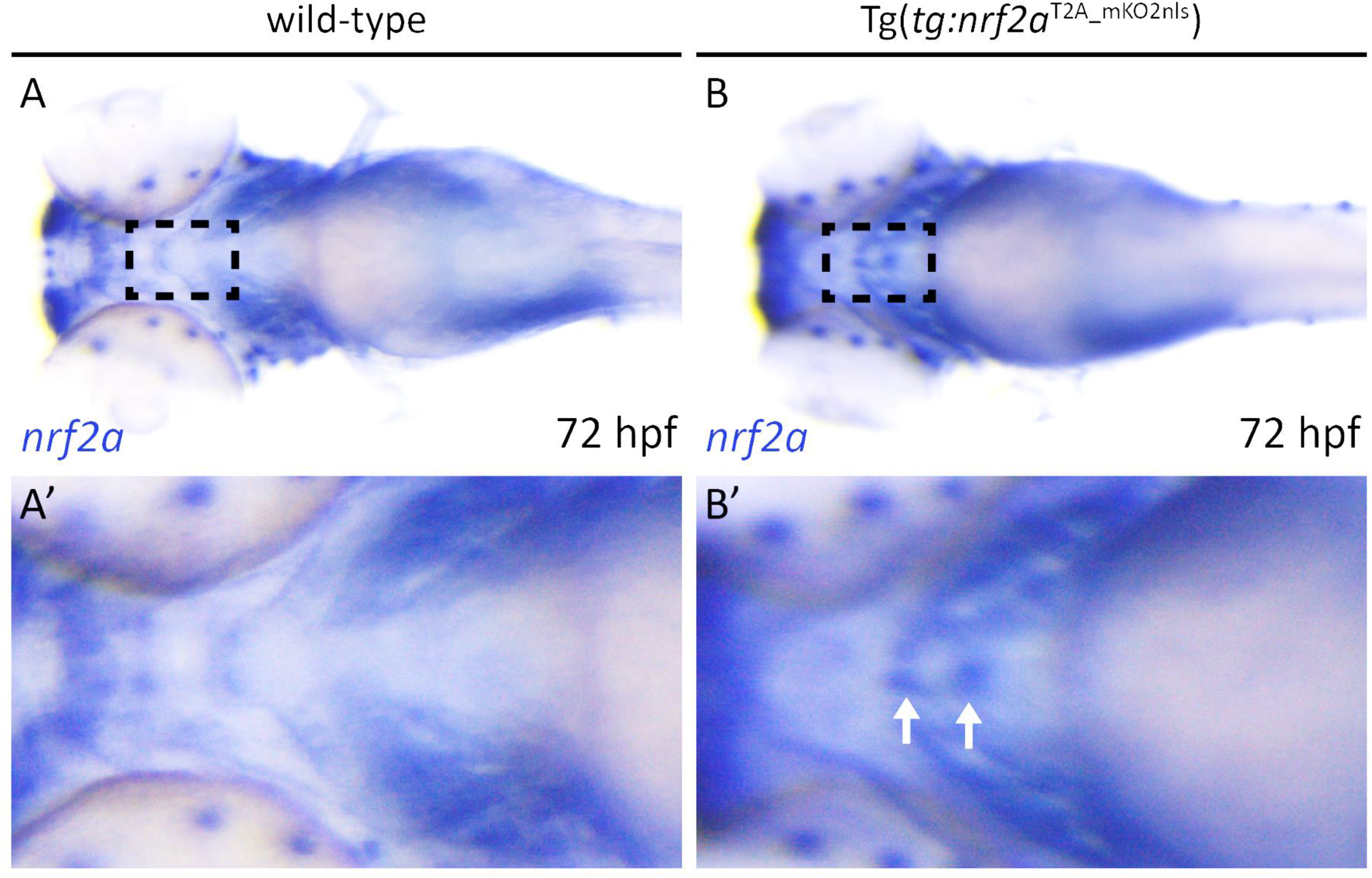
Tg(*tg:nrf2a^T2A_mKO2nls^*) embryos over-express *nrf2a* mRNA in their thyroid gland. (A-B) Detection of *nrf2a* mRNA using whole mount *in situ* hybridisation on 72hpf embryos. Analysis of *nrf2a* expression pattern shows presence of *nrf2a* mRNA in a location corresponding to the developing thyroid gland in Tg(*tg:nrf2a^T2A_mKO2nls^*) embryos but not in wild-type embryos. (A’-B’) show higher magnification of the thyroid region (dashed line square in picture A and B) of the corresponding embryo. White arrows highlight the *nrf2a* mRNA in the developing thyroid. All images are ventral views, anterior to the left.

**Supplementary Figure 4:**
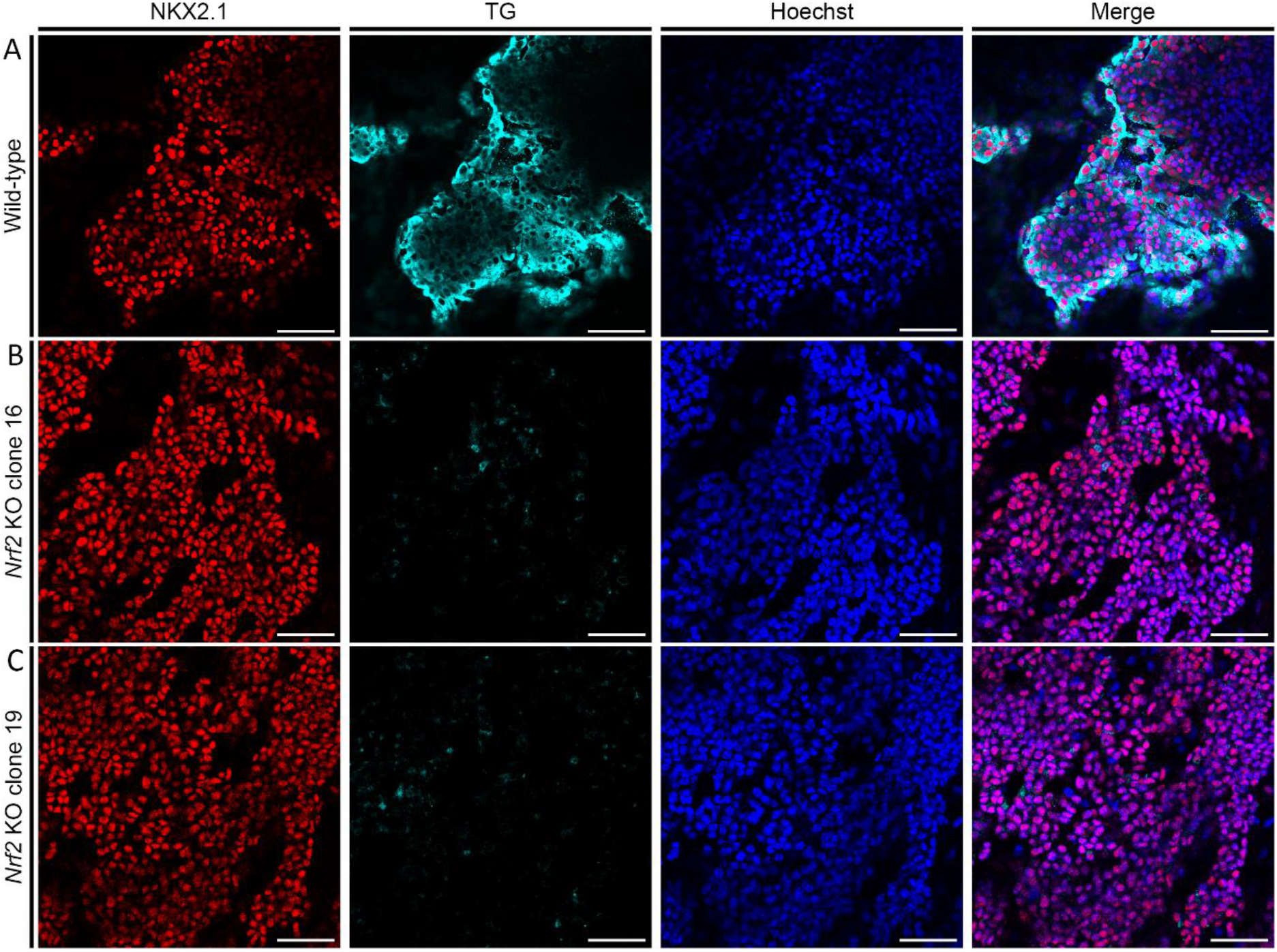
Initial characterisation of Nrf2 KO thyroid organoids. Immunofluorescence experiments revealed that loss of Nrf2 function impairs thyroid differentiation in both Nrf2 KO clones (A-C). (A) Confocal images from Nrf2 wild-type cells at day 22 displaying differentiated organoids marked with NKX2.1 (Red) and TG (Cyan), nuclei were labelled using Hoechst (Blue). (B-C) Confocal images of Nrf2 KO clone 16 (B) and Nrf2 KO clone 19 (C) labelled as described for (A). Scale bars: 100 μm.

**Supplementary Figure 5:**
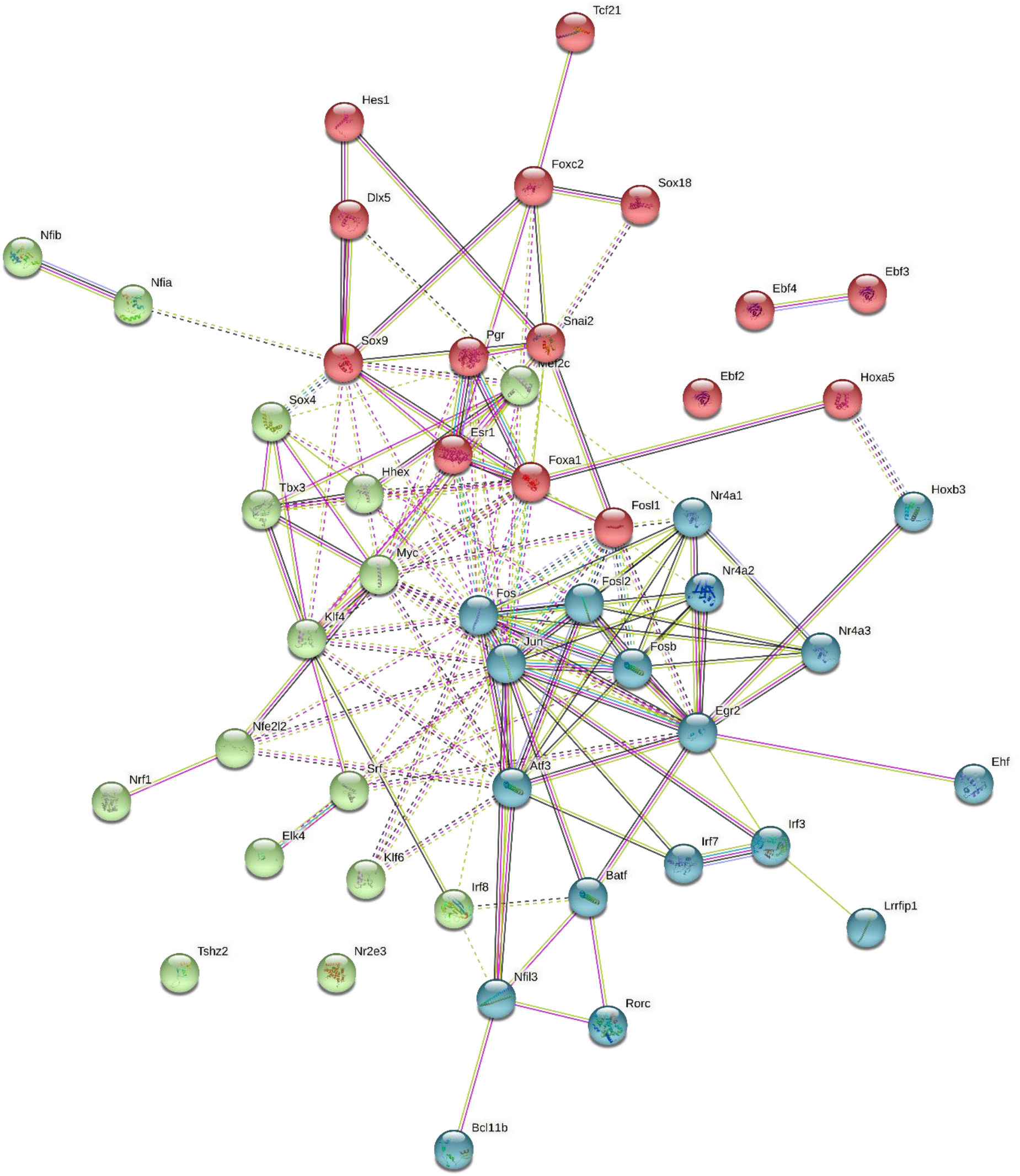
STRING interaction. STRING analysis predicts a strong association network involving the upregulated transcription factors observed in Nrf2 KO condition, with a main core comprising *Fos* and *Jun* genes.

**Supplementary Figure 6:**
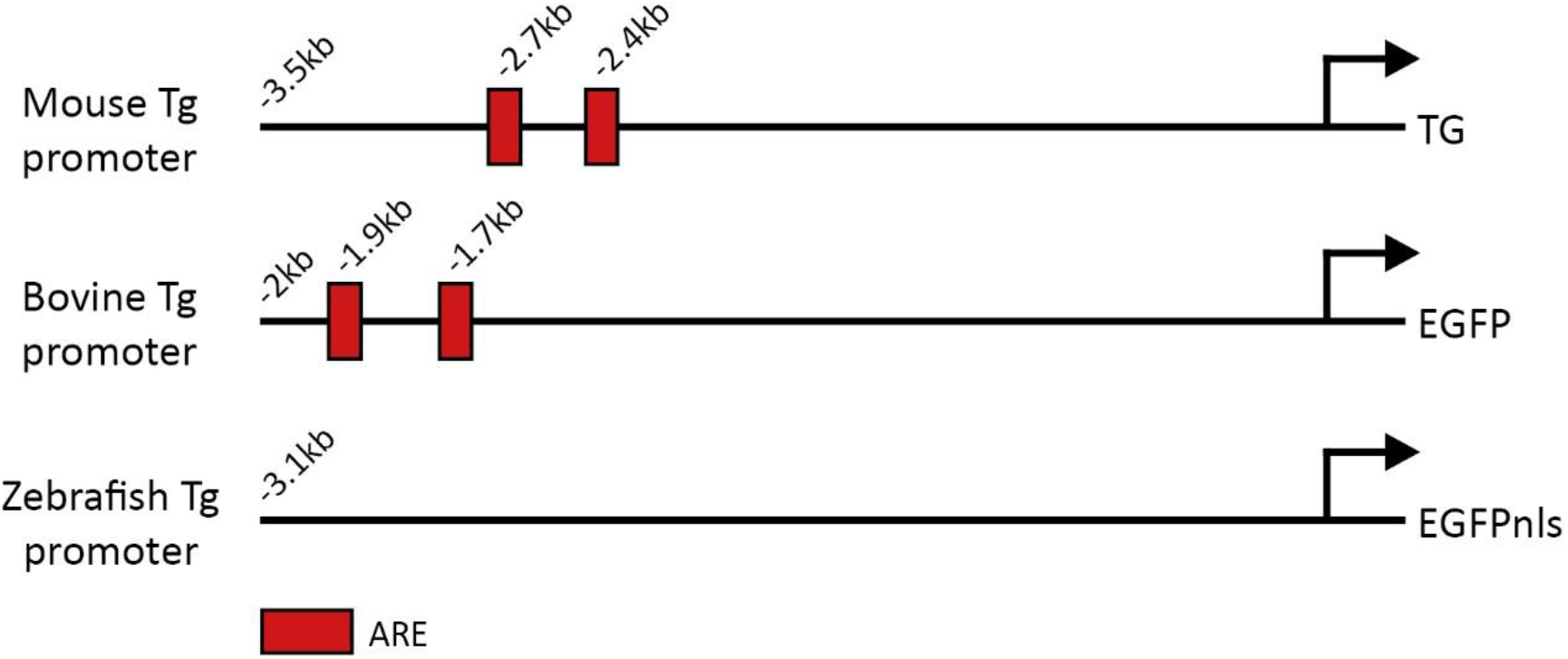
Thyroglobulin promoters analysis. Analysis of the mouse, bovine and zebrafish thyroglobulin promoter for the presence of ARE revealed that only mouse and bovine Tg promoters displays ARE in their sequence. Schematic representation of mouse, bovine and zebrafish Tg promoter based on available sequence (mouse promoter) and sequences used as reporter drivers (bovine and zebrafish promoters). ARE are represented as red boxes and their relative position to the end of the promoters are given.

